# Cytosolic peptide accumulation activates the NLRP1 and CARD8 inflammasomes

**DOI:** 10.1101/2022.03.22.485298

**Authors:** Elizabeth L. Orth-He, Hsin-Che Huang, Sahana D. Rao, Qinghui Wang, Qifeng Chen, Claire M. O’Mara, Ashley J. Chui, Michelle Saoi, Andrew R. Griswold, Abir Bhattacharjee, Daniel P. Ball, Justin R. Cross, Daniel A. Bachovchin

**Author notes:** Correspondence to Daniel A. Bachovchin. These authors contributed equally (Elizabeth L. Orth-He, Hsin-Che Huang).

## Abstract

NLRP1 and CARD8 are related sensors that form inflammasomes, but the danger signals that they detect are not fully established. These proteins undergo autoproteolysis, generating repressive N-terminal (NT) and inflammatory C-terminal (CT) fragments. The proteasome-mediated degradation of the NT releases the CT from autoinhibition, but the CT is then sequestered in a complex with the full-length sensor and DPP9. Here, we show that cytosolic peptide accumulation activates these inflammasomes. We found that a diverse array of peptides accelerates NT degradation, and those with N-terminal XP sequences also destabilize the ternary complexes. Peptides interfere with many biological processes, including protein folding. We show that unrelated agents that disrupt protein folding also induce NT degradation, but do not cause inflammasome activation because DPP9 sequesters the CT fragments in the absence of XP peptides. Overall, these results indicate that NLRP1 and CARD8 detect protein misfolding that is associated with peptide accumulation.

## INTRODUCTION

Mammals express at least six pattern recognition receptors (PRRs) that sense danger-associated signals and nucleate the formation of signaling platforms called inflammasomes (Broz and Dixit, 2016). Inflammasomes recruit and activate the cysteine protease caspase-1, which then cleaves and activates the pore-forming protein gasdermin D (GSDMD) and (in most cases) the inflammatory cytokines interleukin-1β and -18 (IL-1β/18), triggering a lytic form of cell death called pyroptosis. NLRP1 (nucleotide-binding domain leucine-rich repeat pyrin domain-containing 1) and CARD8 (caspase activation and recruitment domain-containing 8) are related human PRRs that form inflammasomes, and a series of recent studies have made considerable progress in elucidating their activation mechanisms (Chui et al., 2019; Hollingsworth et al., 2021; Huang et al., 2021; Johnson et al., 2018; Okondo et al., 2017; Okondo et al., 2018; Sharif et al., 2021; Zhong et al., 2018). However, the danger signals that these inflammasomes evolved to sense have not yet been definitively established (Bachovchin, 2021).

The human NLRP1 (hNLRP1) protein has an N-terminal pyrin domain (PYD) and a disordered region preceding the nucleotide-binding (NACHT), leucine-rich repeat (LRR), function-to-find (FIIND), and caspase activation and recruitment domains (CARDs) (**Fig. 1A**). CARD8 has an N-terminal disordered stretch of ∼160 amino acids followed by a similar FIIND-CARD region (**Fig. 1A**). The NLRP1 and CARD8 FIINDs undergo autoproteolysis between their ZU5 (ZO-1 and UNC5) and UPA (conserved in UNC5, PIDD, and ankyrin) subdomains, generating N-terminal (NT) and C-terminal (CT) polypeptide chains that remain non-covalently associated in an autoinhibited state (D’Osualdo et al., 2011; Finger et al., 2012; Frew et al., 2012). Mice have two functional NLRP1 homologs (mNLRP1A and B) and rats have one NLRP1 homolog (rNLRP1), but neither rodent species has a CARD8 homolog. The rodent NLRP1 proteins lack the N-terminal PYD but otherwise have domain organizations like hNLRP1. NLRP1 is highly polymorphic both within and across species, and at least five distinct alleles of rNLRP1 and mNLRP1B are present in inbred rat and mouse strains (Boyden and Dietrich, 2006; Newman et al., 2010). Despite their considerable diversity in primary sequence, all NLRP1 and CARD8 alleles are activated via a process called “functional degradation”, in which the proteasome-mediated destruction of the NT fragment releases the CT fragment to nucleate an inflammasome (Chui et al., 2019; Sandstrom et al., 2019).

**Figure 1.**
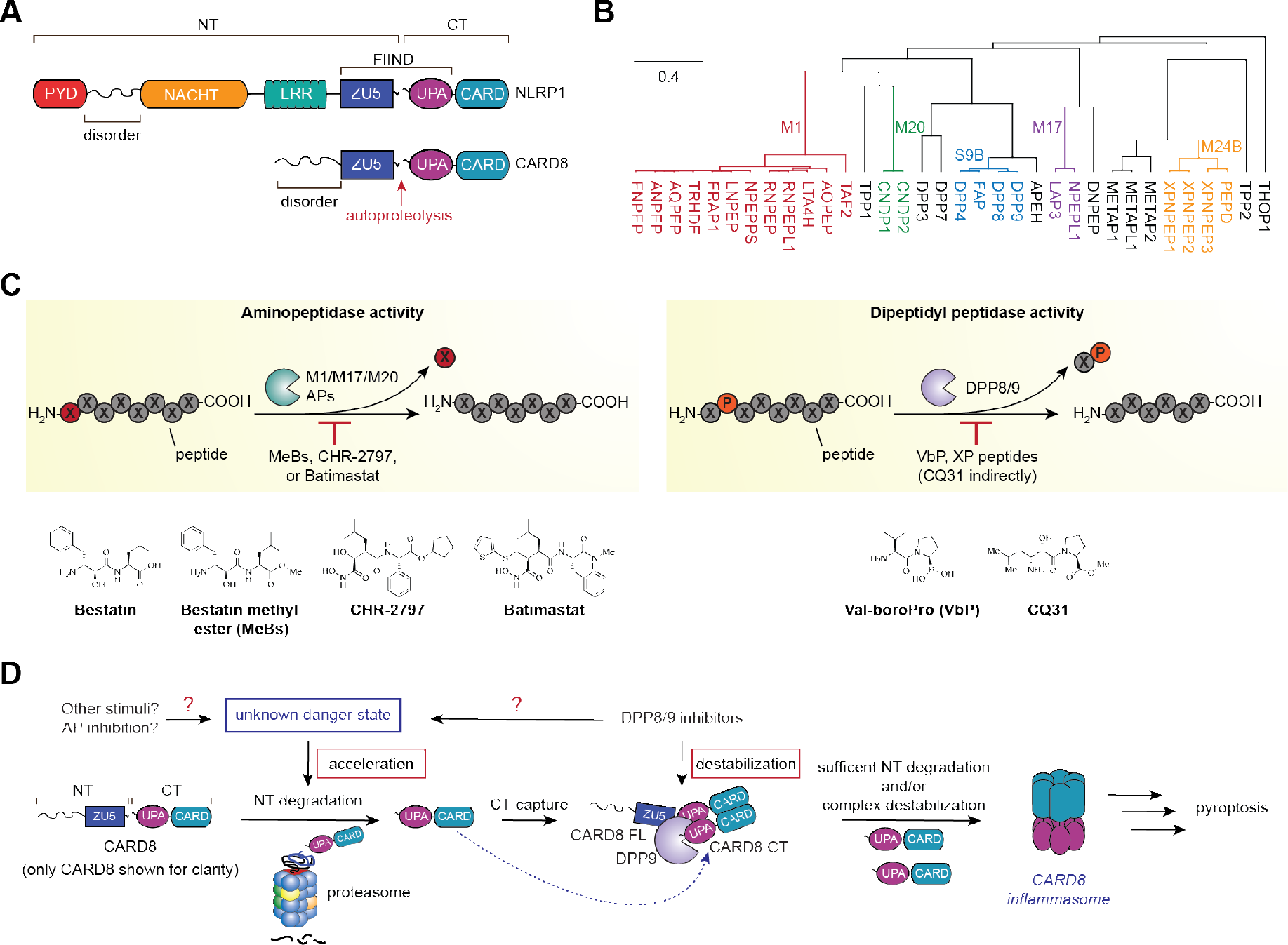
Overview of NLRP1 and CARD8 regulation. (**A**) Domain architecture of human NLRP1 and CARD8. The autoproteolysis sites are indicated. (**B**) Dendrogram of human APs and DPPs involved in peptide hydrolysis. Certain enzyme families are indicated. (**C**) Schematic of bestatin-sensitive AP activity (left) and VbP-sensitive DPP activity (right). Chemical structures of peptidase inhibitors used in this study are shown. (**D**) Proposed model for NLRP1 and CARD8 inflammasome activation. Constitutive low-level proteasome-mediated degradation of the CARD8/NLRP1 NT fragment releases the CT fragment, which is then captured as part of a ternary complex that also includes DPP9 and full-length (FL) CARD8/NLRP1. Acceleration of NT degradation or destabilization of the ternary complex enables enough CT fragments to escape repression and oligomerize into an inflammasome. DPP8/9 inhibitors, and potentially other stimuli, might induce a danger state that accelerates NT degradation. DPP8/9 inhibitors also destabilize the ternary complexes. See also Tables S1 and S2.

A wide array of unrelated pathogen-derived stimuli, including bacterial and viral proteases, E3 ligases, and long dsRNA, have been reported to activate at least one human or rodent NLRP1 allele via functional degradation (**Table S1**) (Boyden and Dietrich, 2006; Hornung et al., 2009; Robinson et al., 2020; Sandstrom et al., 2019; Tsu et al., 2020). The most well-studied of these is the anthrax lethal factor (LF) metalloprotease, which cleaves some rodent NLRP1 alleles near their N-termini and thereby generates unstable neo-N-termini that are rapidly recognized and degraded by the N-end rule proteasome degradation pathway (Chui et al., 2019; Frew et al., 2012; Sandstrom et al., 2019). Notably, none of these pathogen-derived stimuli, including LF, universally activate all functional human and rodent NLRP1 alleles. Similarly, HIV protease recently was discovered to cleave human CARD8 and thereby activate the CARD8 inflammasome, but not the rodent or human NLRP1 proteins (Wang et al., 2021). Collectively, these observations have fueled speculation that the various polymorphic human and rodent NLRP1 alleles and human CARD8 each evolved independently to detect one or more entirely distinct pathogen-associated signals.

Alternatively, it is possible that all NLRP1 alleles (and potentially CARD8) evolved to sense a single, specific cellular danger state that has yet to be identified (Bachovchin, 2021). Consistent with this premise, potent inhibitors of the dipeptidyl peptidases 8 and 9 (DPP8/9) activate all functional NLRP1 and CARD8 alleles (**Table S1**) (Gai et al., 2019; Johnson et al., 2018; Okondo et al., 2018; Zhong et al., 2018). DPP8/9 are serine proteases that cleave XP dipeptides (X is any amino acid) from the N-termini of unstructured polypeptides (**Fig. 1B,C**) (Geiss-Friedlander et al., 2009; Griswold et al., 2019b; Tang et al., 2009). Potent DPP8/9 inhibitors, including Val-boroPro (VbP, **Fig. 1C**), induce inflammasome formation via two distinct mechanisms. First, the inhibition of DPP8/9’s enzymatic activity accelerates the proteasome-mediated degradation of many misfolded and disordered proteins, including the CARD8^NT^ and NLRP1^NT^ fragments, via a poorly understood pathway (**Fig. 1D**) (Chui et al., 2020; Chui et al., 2019; Griswold et al., 2019a). Second, potent DPP8/9 inhibitors destabilize a repressive ternary complex that forms between DPP9 (or DPP8), the full-length (FL) PRR, and the liberated CT fragment that restrains spurious inflammasome formation by low levels of the free CT (**Fig. 1D**) (Hollingsworth et al., 2021; Huang et al., 2021; Sharif et al., 2021). Thus, DPP8/9 appear to be somehow connected to the primordial function of NLRP1 and CARD8 inflammasomes, including to a specific danger state that these inflammasomes universally sense (**Fig. 1D**) (Bachovchin, 2021). However, the identity of the danger state linked to DPP8/9 remains unknown.

Bestatin and bestatin methyl ester (MeBs, a more cell permeable analog of bestatin) are non-specific inhibitors of M1, M17, and M20 metallo-aminopeptidases (APs), which are enzymes that cleave single N-terminal amino acids from polypeptide chains (**Fig. 1B,C**) (Burley et al., 1991; Suda et al., 1976; Tsuge et al., 1994). MeBs has been reported to elicit several biological effects, including blockade of the N-end rule pathway (Wickliffe et al., 2008), degradation of the cellular inhibitor of apoptosis protein 1 (cIAP1) (Sekine et al., 2008), and starvation of intracellular amino acids (Krige et al., 2008; Vabulas and Hartl, 2005). Intriguingly, we found that MeBs induces pyroptotic cell death in *DPP8^−/−^/DPP9^−/−^* (*DPP8/9^−/−^*), but not in wild-type, THP-1 cells (Chui et al., 2019). MeBs also synergizes with VbP to induce more pyroptosis in THP-1 cells, RAW 264.7 cells, and primary bone-marrow derived macrophages (BMDMs) from Sprague-Dawley (SD) rats (Chui et al., 2019; Gai et al., 2019) as well as more serum G-CSF production in C57BL6/J mice (Chui et al., 2019). As these cell types and animals all express CARD8 and NLRP1 alleles (**Table S2**), MeBs appears to augment the activation of all NLRP1 and CARD8 alleles after genetic or pharmacologic inactivation of DPP8/9. Notably, the structurally unrelated AP inhibitors CHR-2797 and batimastat (**Fig. 1C**) similarly induce synergistic pyroptosis, strongly suggesting these responses are due to AP inhibition and not some off-target activity of MeBs (Chui et al., 2019).

Thus, it appears that AP inhibition also contributes in some way to the inflammasome-activating danger state (**Fig. 1D**).

Here, we investigated the mechanistic basis for the synergy between AP and DPP8/9 inhibitors. We found that AP inhibition alone (i.e., without simultaneous DPP8/9 inhibition) induces the accumulation of many proteasome-derived peptides, but not those with XP N-termini, and thereby accelerates the degradation of the NT fragments. However, DPP8/9 ternary complexes effectively quench the CT fragments released by this mechanism. Peptides are known to interfere with many biological processes, including chaperone-mediated protein folding (Li et al., 2003; Otvos et al., 2000). Notably, we found that distinct agents that disrupt protein folding similarly stimulate NT degradation, but pyroptosis is again suppressed by the DPP8/9 complexes. We show that proteasome-derived peptides with XP N-termini, which are mimicked by VbP, must also accumulate to overcome the DPP8/9 checkpoint. Overall, we propose that NLRP1 and CARD8 detect protein misfolding linked to cytosolic peptide build-up.

## RESULTS

### AP inhibitors synergize with DPP8/9 inhibitors

We first wanted to comprehensively characterize the relationship between AP inhibitors and inflammasome activation. VbP activates the CARD8 inflammasome in several human acute myeloid leukemia cell lines, including THP-1 (albeit only minimally after 6 h), MV4;11, and OCI- AML2 cells (**Table S2**) (Johnson et al., 2018). As expected, we found that VbP induced at least some pyroptosis in these cells after 6 h, as evidenced by LDH release and GSDMD cleavage (into the p30 NT fragment) assays (**Fig. 2A–C, Fig. S1A–B**). Also as expected, VbP did not induce any additional pyroptosis in *DPP8/9^−/−^* THP-1 cells (**Fig. 2A**). In contrast and as previously reported, MeBs, CHR-2797, and batimastat did not induce pyroptosis in wild-type (WT) THP-1 cells, but did induce pyroptosis in *DPP8/9^−/−^* THP-1 cells (**Fig. 2A**) and in WT THP-1, MV4;11, and OCI-AML2 cells also treated with VbP (**Fig. 2B,C, Fig. S1A,B**) (Chui et al., 2019). This death was entirely mediated by CARD8, as *CARD8^−/−^* cells were completely resistant to the combinations (**Fig. 2B, Fig. S1B**). Similarly, MeBs also induced synergistic pyroptosis with VbP in human naive CD3 T cells (**Fig. 2D**), which express a functional CARD8 inflammasome (Johnson et al., 2020; Linder et al., 2020). Thus, AP inhibitors increase VbP- and *DPP8/9* knockout-induced inflammasome activation in both immortalized and primary cells with endogenous CARD8. It should be noted that these AP inhibitors were not toxic on their own at these 6 h time points, but that they do induce apoptosis over longer (≥ 12 h) intervals, as evidenced by caspase-1-independent death involving poly(ADP-ribose) polymerase (PARP) cleavage (**Fig. S1C,D**).

**Figure 2.**
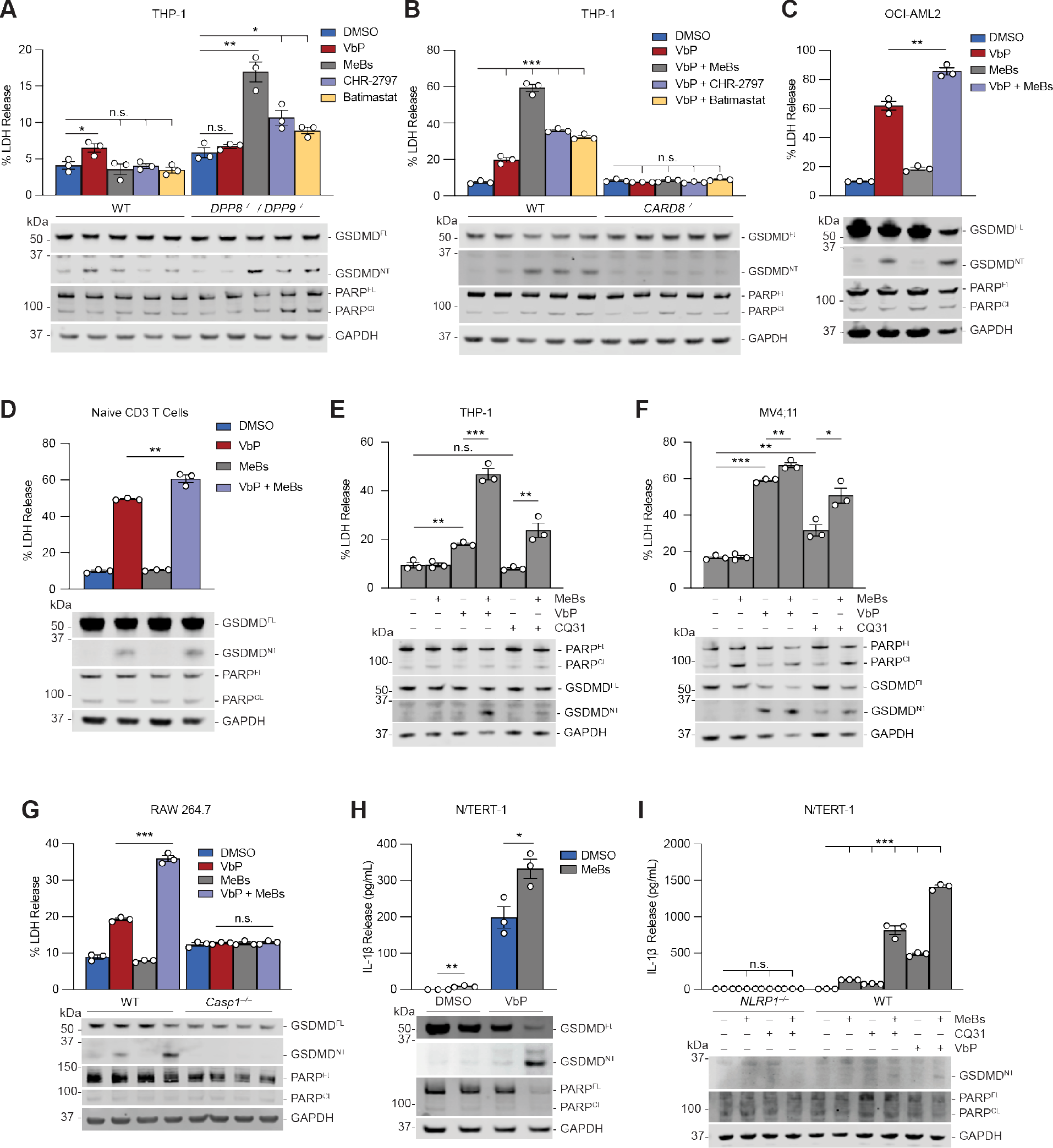
Metallo-aminopeptidase inhibitors induce synergistic inflammasome activation with DPP8/9 blockade. (**A,B**) The indicated THP-1 cells were treated with VbP (10 µM), MeBs (10 µM), CHR-2797 (10 µM), or batimastat (10 µM) for 6 h before LDH release and immunoblot analyses. (**C,D**) OCI-AML2 or resting human CD3 T cells were treated with VbP (10 µM) and/or MeBs (10 µM) for (**C**) 6 h or (**D**) 18 h before LDH release and immunoblot analyses. (**E,F**) THP-1 or MV4;11 cells were treated with VbP (10 µM), CQ31 (20 µM), and/or MeBs (10 µM) for 16 h before LDH release and immunoblot analyses. (**G**) WT or *Casp1^−/−^* RAW 264.7 cells were treated with VbP (10 µM) and/or MeBs (10 µM) for 6 h before LDH release and immunoblot analyses. (**H,I**) WT or *NLRP1^−/−^* human N/TERT-1 keratinocytes were treated with VbP (0.2 µM), MeBs (20 µM), or CQ31 (20 µM) for 24 h before IL-1β release and immunoblot analyses. Data are means ± SEM of 3 replicates. All data, including immunoblots, are representative of three or more independent experiments. *** *p* < 0.001, ** *p* < 0.01, * *p* < 0.05 by two-sided Students *t*-test. n.s., not significant. See also Figure S1.

We recently discovered that weak inhibitors of DPP8/9 (e.g., those with IC50 values > 10 µM, ∼ 4 orders of magnitude higher than VbP), including Val-Pro (VP) and Ile-Pro (IP) dipeptides, selectively activate the CARD8 inflammasome (Rao et al., 2022). These molecules do not activate NLRP1 likely because the NLRP1^CT^, unlike the CARD8^CT^, directly contacts the DPP8/9 active site in the ternary complex and forms a tighter interaction that is more difficult to sufficiently destabilize (Hollingsworth et al., 2021; Sharif et al., 2021). High concentrations of these dipeptides can be introduced into cells in three ways: 1) treatment with cell permeable esterified dipeptides, including VP methyl ester (VP-OMe); 2) treatment with the small molecule CQ31 (**Fig. 1C**), which inhibits the M24B aminopeptidases prolidase (PEPD) and Xaa-Pro aminopeptidase 1 (XPNPEP1) and blocks the hydrolysis of endogenous XP dipeptides (Rao et al., 2022); or 3) by genetic knockout of *PEPD* and/or *XPNPEP1*. We found that MeBs synergizes with VP-OMe and CQ31 to induce more CARD8-dependent pyroptosis (**Fig. 2E,F, Fig. S1E,F**). Similarly, we found that MeBs induced pyroptosis in *PEPD^−/−^* THP-1 cells, *PEPD^−/−^/ XPNPEP1^−/−^* THP-1 cells, and *PEPD^−/−^* MV4;11 cells (**Fig. S1G,H**), as well as in *PEPD^−/−^* HEK 293T cells ectopically expressing the CARD8 inflammasome components (**Fig. S1I**). In contrast, knockout of *XPNPEP1* alone did not engender MeBs sensitivity, demonstrating that PEPD is the more important of these two enzymes in restraining CARD8 activation (**Fig. S1H**). Overall, these data show that MeBs also synergizes with XP-containing peptides to more strongly activate the CARD8 inflammasome.

We next wanted to investigate the impact of MeBs on NLRP1 inflammasome activation. We found that MeBs induced synergistic pyroptosis with VbP in RAW 264.7 and J774.1 mouse macrophages (**Fig. 2G**, **Fig. S1J**), both of which express mNLRP1B allele 1 (**Table S2**), and in immortalized human N/TERT-1 keratinocytes (**Fig. 2H**), which express hNLRP1 (**Table S2**). It should be noted that the human NLRP1 inflammasome, but not the human CARD8 inflammasome, releases cleaved IL-1β (Ball et al., 2020), and therefore we evaluated IL-1β release in N/TERT-1 keratinocytes. In addition, MeBs induced substantial NLRP1-dependent ASC speck formation in *DPP9^−/−^*, but not control, HEK 293T cells ectopically expressing human NLRP1 and GFP-tagged ASC (**Fig. S1K**). As mentioned above, CQ31 does not activate the human or rodent NLRP1 inflammasome as a single agent (Rao et al., 2022). Intriguingly, however, we found that the combination of CQ31 and MeBs does activate NLRP1 in N/TERT-1 keratinocytes (**Fig. 2I**). In contrast, this drug combination still did not activate the mouse NLRP1B in RAW 264.7 cells (**Fig. S1L**), which we speculate is because the mouse NLRP1 inflammasomes have particularly high activation thresholds (Cirelli et al., 2014; Ewald et al., 2014; Johnson et al., 2020). Regardless, these data, coupled with our previous results (Chui et al., 2019), show that AP inhibitors synergize with VbP, XP peptides, or *DPP8/9* knockout to induce greater CARD8 and NLRP1 inflammasome responses. Importantly, MeBs did not impact lipopolysaccharide (LPS) plus nigericin activation of the NLRP3 inflammasome in THP-1 cells (**Fig. S1M**), showing that this synergy is specific to the NLRP1 and CARD8 inflammasome pathways.

### MeBs accelerates NT degradation

We next wanted to determine if AP inhibitors induce synergistic inflammasome activation by accelerating NT degradation or by destabilizing the DPP8/9 ternary complexes (**Fig. 1D**). As MeBs does not interact with DPP8/9 (Rao et al., 2022), we reasoned that MeBs likely does not directly interfere with the ternary complexes. Indeed, we previously found that MeBs does not displace NLRP1 or CARD8 from immobilized DPP9 *in vitro* (Hollingsworth et al., 2021; Sharif et al., 2021). Moreover, we demonstrated above that AP inhibitors induce pyroptosis in *DPP8/9^−/−^* THP-1 cells that completely lack these repressive complexes (**Fig. 2A**). However, it remained possible that AP inhibition indirectly destabilizes the ternary complexes in cells, and that this, at least in part, contributes to inflammasome activation. We previously developed a method to test ternary complex destabilization in cells that leverages the degradation tag (dTAG) system, in which the small molecule dTAG-13 is used to rapidly trigger the degradation of proteins with FKBP12^F36V^ tags (dTAGs) (**Fig. 3A**) (Nabet et al., 2018; Sharif et al., 2021). Briefly, we create the DPP8/9 ternary complex in cells by ectopically expressing the CARD8 <u>Z</u>U5-<u>U</u>PA-<u>C</u>ARD region with an N-terminal dTAG (dTAG-CARD8^ZUC^) together with an autoproteolysis-defective S297A mutant CARD8 FIIND domain (FIIND^SA^); treatment of these cells with dTAG-13 releases the free CARD8^CT^ to form ternary complexes with the FIIND^SA^ and endogenous DPP9 (**Fig. 3A**). DPP8/9 inhibitors, including VbP, compound 8j, and VP-OMe, destabilize these complexes and induce pyroptosis in HEK 293T cells that also stably express CASP1 and GSDMD (**Fig. 3A, Fig. S2A**). It should be noted that DPP8/9 inhibitors do not activate dTAG-CARD8^ZUC^ on their own (i.e., without dTAG-13) because this fusion protein lacks the N-terminal disordered region of CARD8 required for DPP8/9 inhibitor-induced pyroptosis (Chui et al., 2020). Notably, we found that MeBs did not induce additional cell death on its own or in combination with VbP after dTAG-13 treatment (**Fig. 3B, Fig. S2A**). Thus, MeBs does not directly or indirectly destabilize the repressive DPP8/9 ternary complexes, at least to the extent we can determine using the assays described above.

**Figure 3.**
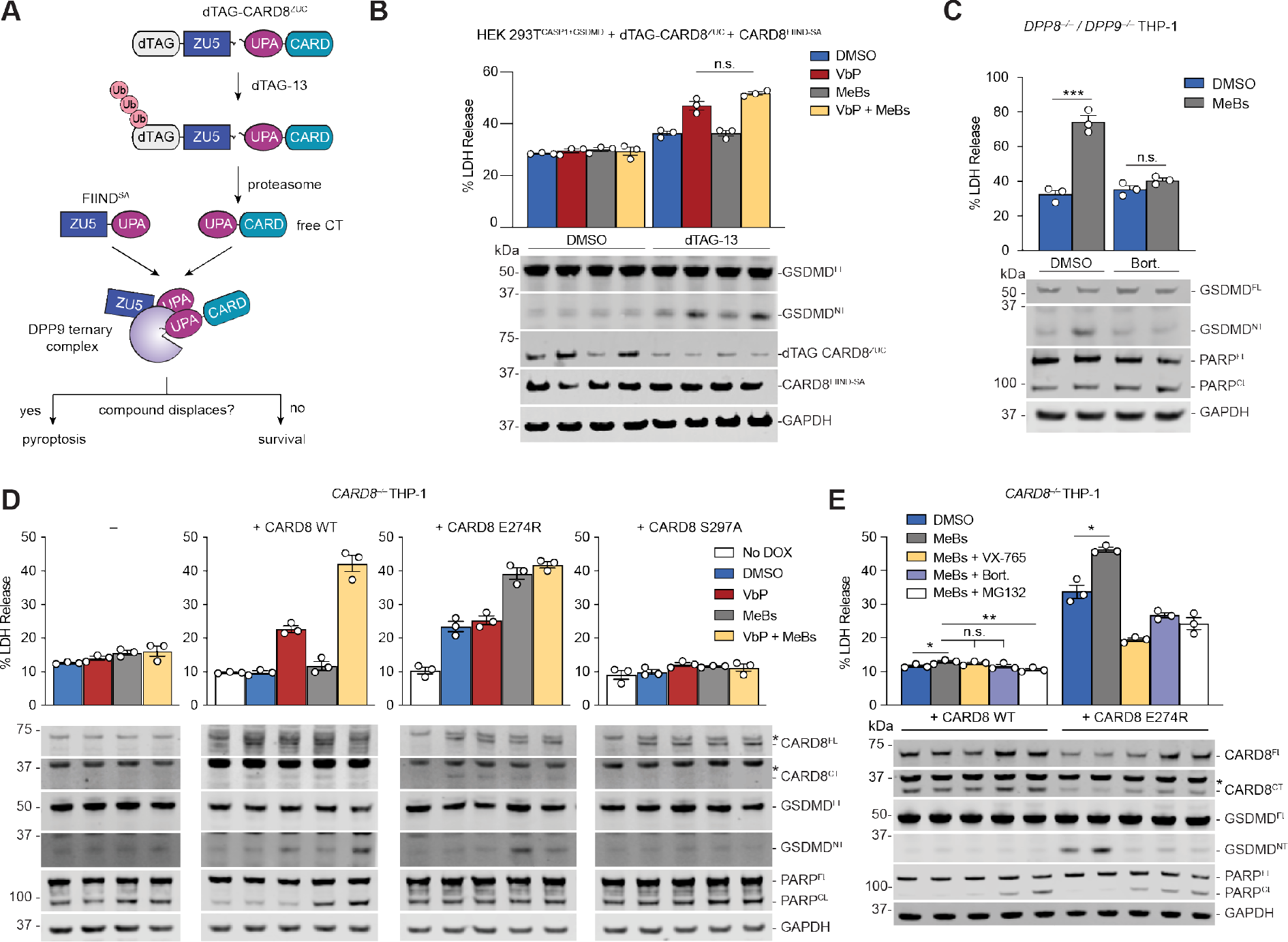
AP inhibitors accelerate CARD8^NT^ degradation. (**A**) Schematic of the experiment to assess DPP9-CARD8 ternary complex displacement in cells. (**B**) HEK 293T^CASP1+GSDMD^ cells were transiently transfected plasmids encoding dTAG-CARD8^ZUC^ and the isolated FIIND^SA^ 48 h prior to treatment with dTAG-13 (500 nM), VbP (10 µM), MeBs (10 µM), or the combination for 6 h. Samples were collected and analyzed by immunoblotting and LDH release. (**C**) *DPP8^−/−^/DPP9^−/−^* THP-1 cells were treated with MeBs (10 µM) and/or bortezomib (Bort., 10 µM) for 8 h before LDH and immunoblot analyses. (**D,E**) *CARD8^−/−^* THP-1 cells stably containing doxycycline (DOX)- inducible CARD8 WT, E274R, and/or S297A were induced with 100 ng/mL DOX for 16 h prior to treatment with VbP (10 µM), MeBs (10 µM), VX765 (50 µM), Bort. (10 µM), and/or MG132 (10 µM) for 6 h. Samples were then collected for LDH release and immunoblot analyses. Note in (**D**) that immunoblot and LDH analyses were performed separately. Data are means ± SEM of 3 replicates. All data, including immunoblots, are representative of three or more independent experiments. *** *p* < 0.001, ** *p* < 0.01, * *p* < 0.05 by two-sided Students *t*-test. n.s., not significant.See also Figure S2.

Instead, we hypothesized that MeBs mainly accelerated NT degradation. We found that neither VbP, MeBs, nor the combination induced visible CARD8^FL^ or CARD8^NT^ depletion in *CASP1^−/−^* MV4;11 cells by immunoblotting after 6 h (**Fig. S2B,C**). However, this result was not surprising, as only small amounts of free CTs are needed to form inflammasomes (Chui et al., 2019; Sandstrom et al., 2019). We previously found that VbP does cause some observable CARD8^NT^ depletion over longer treatment times (Chui et al., 2020); MeBs similarly appeared to induce some CARD8^NT^ depletion after 48 h (**Fig. S2D**), but this result was not statistically significant (*p* = 0.059) and was potentially confounded by the toxicity of MeBs over longer incubation times (**Fig. S1C,D**). Nevertheless, we found that the proteasome inhibitor bortezomib completely blocked MeBs-induced pyroptosis in *DPP8/9^−/−^* and *PEPD^−/−^* THP-1 cells (**Fig. 3C**, **Fig. S2E**), as well as MeBs plus VbP-induced pyroptosis in WT MV4;11 cells (**Fig. S2F**). Thus, MeBs likely stimulates synergistic pyroptosis by triggering more NT degradation. We provide additional evidence that AP inhibition indeed induces NT degradation using NLRP1 and CARD8 mutants that cannot bind DPP8/9, as described below.

### The DPP9 ternary complex restrains AP inhibitor-induced pyroptosis

We next wanted to determine why AP inhibitors only activated the NLRP1 and CARD8 inflammasomes in combination with DPP8/9 inhibitors. We envisioned two possibilities: 1) AP inhibition alone accelerates NT degradation, but that the CT fragments freed by this mechanism are effectively quenched by DPP8/9 ternary complexes in the absence of DPP8/9 inhibitors; or 2) AP activity is only critical in the absence of DPP8/9 activity, perhaps because APs cleave pyroptosis-inducing DPP substrates to some extent.

To distinguish between these possibilities, we needed to evaluate the impact of MeBs on inflammasome activation in cells with DPP8/9 enzymatic activity but without the ability to form DPP8/9 ternary complexes. We previously discovered that E274R mutant CARD8 does not bind to DPP9 and could be used for such an experiment (Sharif et al., 2021). We next generated *CARD8^−/−^* THP-1 cells containing doxycycline (DOX)-inducible WT CARD8, DPP9-non-binding CARD8 E274R, or autoproteolysis-dead CARD8 S297A constructs (**Fig. 3D**). As expected, cells ectopically expressing WT CARD8 responded to VbP and MeBs like WT THP-1 cells (see **Fig. 2A**,**B**), whereas cells expressing CARD8 S297A were non-responsive (**Fig. 3D**). Also as expected, the expression of CARD8 E274R caused some spontaneous death because DPP9 cannot physically restrain free CARD8^CT^ fragments generated during homeostatic protein turnover in these cells. Notably, we found that MeBs induced considerably more pyroptosis in CARD8 E274R-expressing cells (**Fig. 3D**,**E**), and that this death was blocked by the CASP1 inhibitor VX765 and the proteasome inhibitors bortezomib and MG132 (**Fig. 3E**). Collectively, these data indicate that MeBs alone (e.g., without VbP co-treatment) accelerates CARD8^NT^ degradation, but that DPP8/9 and CARD8^FL^ effectively sequester the released CARD8^CT^ fragments in ternary complexes. DPP8/9 inhibitors destabilize these repressive complexes and thereby synergize with AP inhibitors. Intriguingly, VbP itself did not cause significantly more death in cells expressing CARD8 E274R (**Fig. 3D**), indicating that VbP is a much weaker inducer of CARD8^NT^ degradation than MeBs.

We next wanted to perform an analogous experiment with NLRP1. We previously established that NLRP1 LL1193EE and NLRP1 P1214R have abolished and weakened DPP9 binding, respectively (Hollingsworth et al., 2021). We therefore generated *CARD8^−/−^* THP-1 cells containing doxycycline (DOX)-inducible WT NLRP1, NLRP1 LL1193EE, NLRP1 P1214R, and NLRP1 S1213A (autoproteolysis-dead) constructs. As expected, cells ectopically expressing WT NLRP1 and NLRP1 S1213A responded to VbP and MeBs like NLRP1 WT and knockout cells, respectively (**Fig. S2G**). We found that the ectopic expression of NLRP1 LL1193EE triggered a high level of spontaneous pyroptosis that was not increased by either VbP or MeBs, suggesting death is already at a maximal level that cannot be increased further. Interestingly, we found that the expression of NLRP1 P1214R induced some spontaneous death, which was further increased by both MeBs and VbP. Bortezomib and MG132 attenuated MeBs-enhanced cell death (**Fig. S2H**), indicating that MeBs was similarly accelerating NLRP1^NT^ degradation. As P1214R mutant NLRP1 still retains some binding with DPP9 (Hollingsworth et al., 2021), VbP’s activity in this assay is likely due, at least in part, to ternary complex disruption. Regardless, these data show that AP inhibition alone induces NLRP1^NT^ degradation, but that the DPP8/9 ternary complex effectively sequesters the inflammasome-forming CT fragments in these cells.

### Amino acid starvation does not activate NLRP1 and CARD8

We next wanted to identify the AP inhibitor-induced “danger signal” that accelerates NT degradation. Notably, the proteasome digests virtually all intracellular proteins at some point into peptides, and then numerous cytosolic peptidases rapidly hydrolyze these peptides into free amino acids (Kisselev et al., 1999). Intriguingly, both M1/17/20 APs and DPP8/9 are thought to be involved in the catabolism of proteasome-generated peptides (**Fig. 1C**, **Fig. 4A**) (Geiss-Friedlander et al., 2009; Griswold et al., 2019b; Saric et al., 2004). As such, we hypothesized that MeBs and VbP might interfere with peptide catabolism, thereby causing cytosolic peptide accumulation and/or amino acid starvation, one of which might trigger inflammasome activation (**Fig. 4A**).

**Figure 4.**
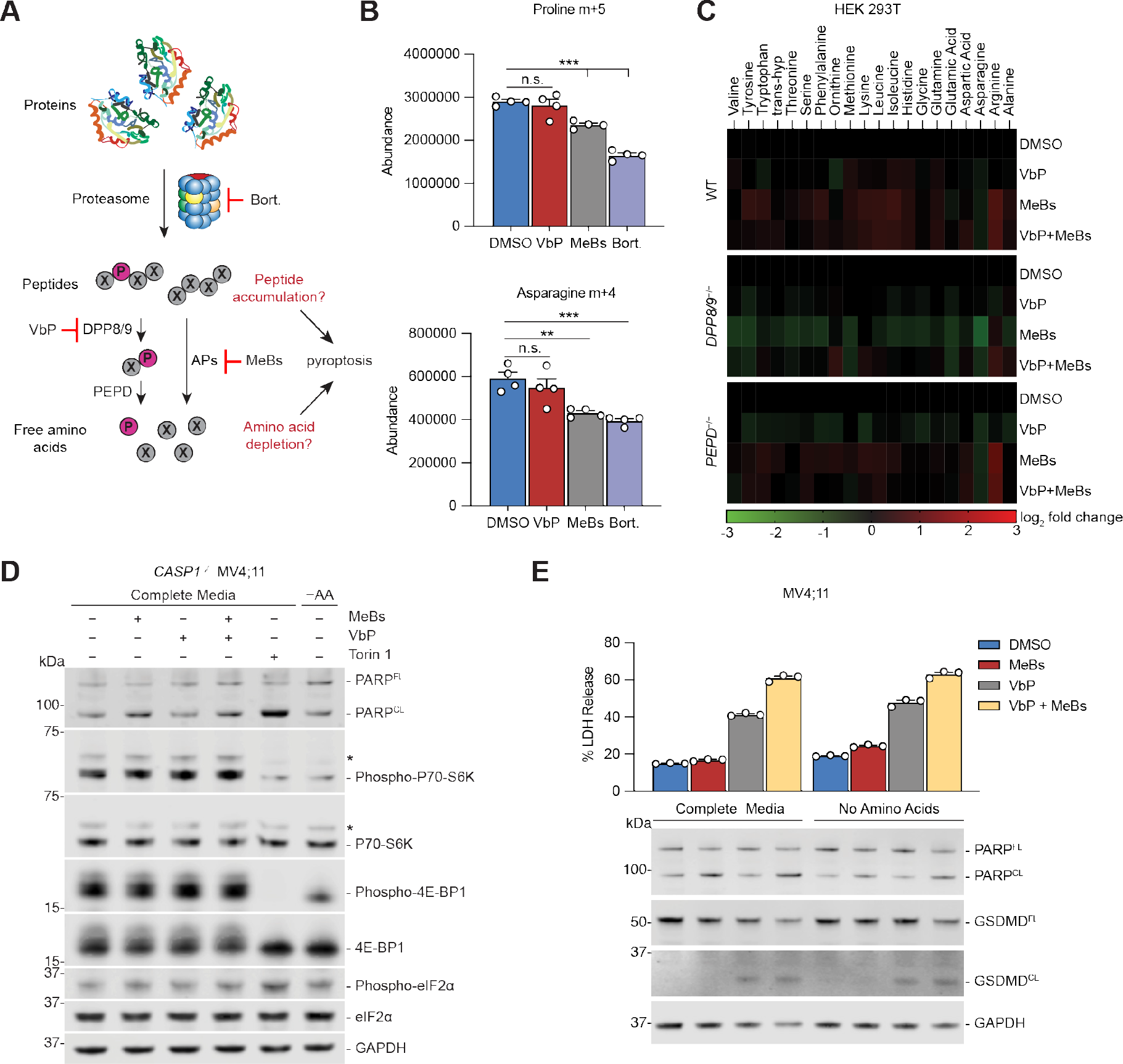
Amino acid depletion does not activate NLRP1 or CARD8. (**A**) Schematic of amino acid recycling from proteins. (**B**) HEK 293T cells were cultured in media supplemented with [U- ^13^C]glutamine and [U-^13^C]leucine for 3 weeks prior to replacement with unlabeled media and simultaneous treatment with VbP (10 µM), MeBs (10 µM) or Bortezomib (Bort., 10 µM). Following 6 h of treatment, cell extracts were profiled for levels of glutamine-derived ^13^C-labeled amino acids. Data are means ± SEM of 4 replicates. (**C**) Heatmap of amino acid levels in WT, *DPP8/9^−/−^* and *PEPD^−/−^* HEK 293T cells treated with VbP (10 µM) and/or MeBs (10 µM) for 3 h. (**D**) *CASP1^−/−^* MV4;11 cells were deprived of amino acids or treated with MeBs (10 µM), VbP (10 µM) or Torin1 (10 µM) for 6 h before immunoblot analysis of amino acid deprivation markers. (**E**) MV4;11 cells were cultured in amino acid-free RPMI supplemented with or without MEM amino acids and treated with MeBs (10 µM) and/or VbP (10 µM) for 6 h, followed by LDH release and immunoblot analysis. Data are means ± SEM of 3 replicates unless otherwise stated. *** *p* < 0.001, ** *p* < 0.01 by two-sided Students *t*-test. n.s., not significant. See also Figure S3 and Table S3.

Proteasome inhibitors and MeBs have previously been reported to cause amino acid deprivation, especially during periods of extracellular nutrient restriction (Krige et al., 2008; Suraweera et al., 2012; Vabulas and Hartl, 2005). We therefore first wanted to investigate the relative impacts of bortezomib, MeBs, and VbP on the recycling of amino acids from protein. To do this, we cultured HEK 293T cells in media containing [U-^13^C]-L-leucine and [U-^13^C]-L-glutamine for 23 days to incorporate isotopically labeled leucine, glutamine, and glutamine-derived amino acids (e.g., asparagine and proline) into proteins (**Fig. S3A**). We then replaced this media with unlabeled media, treated cells with DMSO or inhibitors for 6 h, and extracted and quantified small molecule metabolites (**Fig. S3B**). We found that bortezomib and MeBs both significantly slowed the release of heavy proline and asparagine from proteins (**Fig. 4B**), but that only bortezomib significantly slowed the release of heavy leucine (**Fig. S3C**). VbP did not inhibit the recycling of these amino acids, indicating that DPP8/9 do not play a critical a role in amino acid recycling (**Fig. 4B**, **Fig. S3C**). However, even though MeBs inhibited the release of proline and arginine from protein, neither MeBs, VbP, nor the combination significantly decreased the overall level of any amino acid in WT, *DPP8/9^−/−^* or *PEPD^−/−^* HEK 293T cells (**Fig. 4C**, **Table S3**). Consistently, these drugs did not decrease P70-S6K or 4E-BP-1 phosphorylation or increase eIF2α phosphorylation, key markers of intracellular amino acid starvation (**Fig. 4D**). External amino acid deprivation and treatment with the mammalian target of rapamycin (mTOR) inhibitor Torin 1, both of which modulate these markers, were used as controls in this experiment. Furthermore, we found that removal of all amino acids from the media had no impact on VbP- or VbP plus MeBs-induced CARD8-dependent pyroptosis in MV4;11 cells (**Fig. 4E**). Thus, these data show that DPP8/9 and M1/17/20 AP blockade does not impact intracellular amino acid supply, at least during times of extracellular nutrient sufficiency, and more generally that acute amino acid depletion does not cause NLRP1 or CARD8 inflammasome activation.

### AP inhibition causes proteasome-dependent peptide accumulation

Instead, we reasoned that the accumulation of peptides in the cytosol caused by AP inhibition might be the danger-associated signal (**Fig. 4A**). Cytosolic bestatin-sensitive APs are thought to primarily cleave peptides <10 residues long into their constituent amino acids, removing one residue at a time sequentially from their N termini (Saric et al., 2004) (**Fig. S4A**). When a conformationally-restricted proline residue is in the penultimate position from the N-terminus, however, most APs cannot remove the N-terminal amino acid. Instead, DPP8/9 likely act to remove the N terminal Xaa-Pro dipeptide (Geiss-Friedlander et al., 2009; Griswold et al., 2019b), thereby enabling APs to continue digesting the rest of the polypeptide chain (**Fig. S4A**). Consistent with this mechanism, MeBs, bestatin, CHR-2797 and batimastat, but not VbP, significantly inhibited the cleavage of fluorescent Ala-7-amino-4-methylcoumarin (Ala-AMC) and Leu-AMC reporter substrates in cells (**Fig. 5A,B**); in contrast, VbP, but not MeBs, blocked the cleavage of Ala-Pro-AMC in cells (**Fig. 5C**). Moreover, MeBs significantly slowed the release of alanine from a model PASKYLF peptide in lysates, confirming previous data that bestatin- sensitive enzymes also cleave peptides longer than 2-3 amino acids (**Fig. 5D, Fig. S4B**) (Saric et al., 2004).

**Figure 5.**
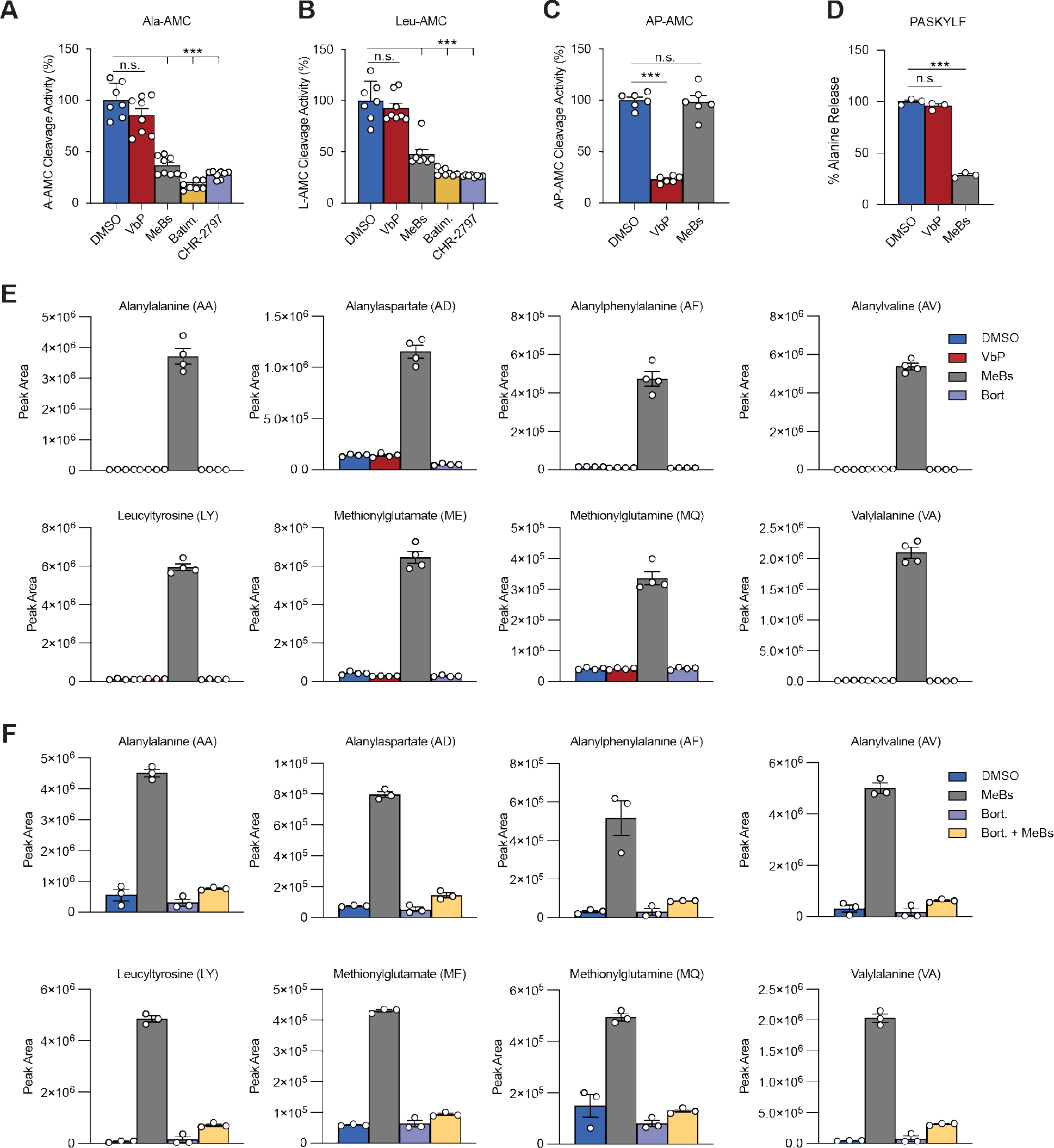
MeBs causes proteasome-derived peptide accumulation. (**A-C**) Inhibition of the indicated peptidase activity in *CARD8^−/−^* THP-1 (**A,B**) or HEK 293T (**C**) cells by VbP (10 µM), MeBs (10 µM), Batimastat (Batim., 10 µM) or CHR-2797 (10 µM). (**D**) HEK 293T cell lysates (0.5 mg/mL) were incubated with the peptide PASKYLF (1 mM) and VbP (10 µM) or MeBs (10 µM) for 6 h prior to quantitation of peptide cleavage by an alanine release assay. (**E**) HEK 293T cells were treated with VbP (10 µM), MeBs (10 µM), or Bortezomib (10 µM) for 6 h. Intracellular metabolites were extracted and the indicated dipeptide concentrations were measured by LC-MS. (**F**) HEK 293T cells were pre-treated with vehicle (DMSO) or Bort. (10 µM) for 30 min, then treated with vehicle (DMSO) or MeBs (10 µM) for 5.5 h. Intracellular metabolites were extracted and dipeptide concentrations were measured by LC-MS. Data are means ± SEM of 3 or more replicates. Peptidase activity data is representative of three or more independent experiments, while endogenous peptide measurement data was collected in a single experiment. *** *p* < 0.001 by two-sided Students *t*-test. n.s., not significant. See also Figure S4.

Based on these results, we predicted that MeBs and VbP would lead to peptide accumulation in cells, whereas bortezomib would conversely deplete peptides. To explore this idea, we next treated cells with either MeBs, VbP, or bortezomib before assessing the levels of various dipeptides, which can be measured by liquid chromatography-mass spectrometry (LC-MS). Consistent with the known instability of cytosolic peptides, we identified very low levels of dipeptides in DMSO- and bortezomib-treated cells (**Fig. 5E**,**F**, **Fig. S4C,D**). However, we observed a striking accumulation of many dipeptides in MeBs-treated HEK 293T and THP-1 cells (**Fig. 5E**,**F, Fig. S4C,D**). Moreover, we found that pre-treatment of cells with bortezomib prior to the addition of MeBs blocked the accumulation of these dipeptides (**Fig. 5F**, **Fig. S4C**), demonstrating that these dipeptides were formed downstream of the proteasome (**Fig. 4A**). We previously showed that CQ31, which inhibits PEPD from cleaving XP dipeptides, causes the accumulation of several Xaa-Pro dipeptides, including GP, IP, and VP, in cells (Rao et al., 2022). Notably, MeBs did not substantially stabilize XP dipeptide levels like CQ31 **(Fig. S4D**), although the level of GP was slightly higher in some MeBs-treated samples (**Fig. S4E**). Thus, MeBs blocks the hydrolysis of many dipeptides, but generally not XPs. These data are consistent with our finding that MeBs does not profoundly destabilize the ternary complexes, although it is possible that the low levels of MeBs-stabilized XP peptides have a small impact on this repressive structure.

Unlike MeBs and CQ31, VbP did not induce the accumulation of any of the dipeptides measured (**Fig. 5E, Fig. S4D,E**), consistent with DPP8/9 only hydrolyzing peptides containing at least three amino acids. Based on the known specificities of DPP8/9 and M1/17/20 APs, we expect that both VbP and MeBs would cause the accumulation of longer proteasome-derived peptides (e.g., see **Fig. 5D**), but, as these longer peptides include a vast array of distinct sequences likely occurring at relatively low concentrations, they are not easily measured using standard metabolomics or proteomics methods. Regardless, these data show that AP inhibition results in the accumulation of proteasome-derived peptides. It should be noted that, because peptides are typically not cell penetrant and any intracellular peptides are rapidly degraded, AP inhibition is to our knowledge the only known way to increase intracellular peptide levels. As such, we are unable to simply introduce high levels of peptides to the cytosol to directly confirm that they accelerate NT degradation.

### cIAP1 is not involved in inflammasome activation

We next sought to investigate potential links between peptide accumulation and accelerated NT degradation. As mentioned above, MeBs is known to induce several additional specific biological effects, including blockade of the N-end rule pathway (Wickliffe et al., 2008) and degradation of the cIAP1 (Sekine et al., 2008), which we reasoned were likely due to the accumulation of intracellular peptides. The N-end rule pathway consists of a number of E3 ligases that bind and ubiquitinate proteins with destabilizing N-terminal residues (Varshavsky, 2011), and we speculate that MeBs induces the accumulation of peptides that compete with destabilizing N-termini for binding to these E3 ligases. However, we previously showed that the N-end rule pathway is not involved in DPP8/9 inhibitor-induced pyroptosis (Chui et al., 2019). Thus, we next turned our attention to cIAP1.

cIAP1 is one of eight mammalian inhibitor of apoptosis proteins (IAPs) involved in controlling caspase activation (Gyrd-Hansen and Meier, 2010; Labbe et al., 2011; Vince et al., 2012). cIAP1 has three N-terminal baculoviral IAP repeat (BIR) domains (BIR1-3) followed by a CARD and a really interesting new gene (RING) finger domain. SMAC and HTRA2, two mitochondrial proteins released into the cytosol during apoptosis, use their first four residues (Ala-Val-Pro-Ile/Ser) called an IAP-binding motif (IBM) to associate with the BIR2 and BIR3 domains of cIAP1 and modulate cIAP1 function (Vaux and Silke, 2003). Most notably, SMAC and HTRA2, as well as small molecule analogs of the IBM, including BV6 and GDC-0152 (Flygare et al., 2012; Varfolomeev et al., 2007), induce the rapid autoubiquitination and degradation of cIAP1. MeBs similarly induces the ubiquitination and degradation of cIAP1 (Sekine et al., 2008), and surface plasmon resonance (SPR), fluorescence polarization, and photoaffinity labeling assays suggested that MeBs directly interacts with cIAP1. However, *in vitro* binding was only observed at much higher concentrations (∼100 µM) than those needed for *in cellulo* cIAP1 degradation (∼3 µM) (Sato et al., 2008; Sekine et al., 2008). We confirmed that MeBs indeed induces proteasome-mediated cIAP1 degradation in cells (**Fig. 6A**, **Fig. S1D, Fig. S5A**). Moreover, we found that the structurally unrelated AP inhibitors CHR-2797 and batimastat also induced cIAP1 degradation, suggesting that this response is likely caused by AP inhibition and not direct cIAP1 binding. Notably, MeBs and batimastat still caused cIAP1 degradation in *SMAC^−/−^*/*HTRA2^−/−^* cells, and thus cIAP1 degradation was not simply due to the activation of these endogenous cIAP1 agonists during apoptosis (**Fig. 6A**).

**Figure 6.**
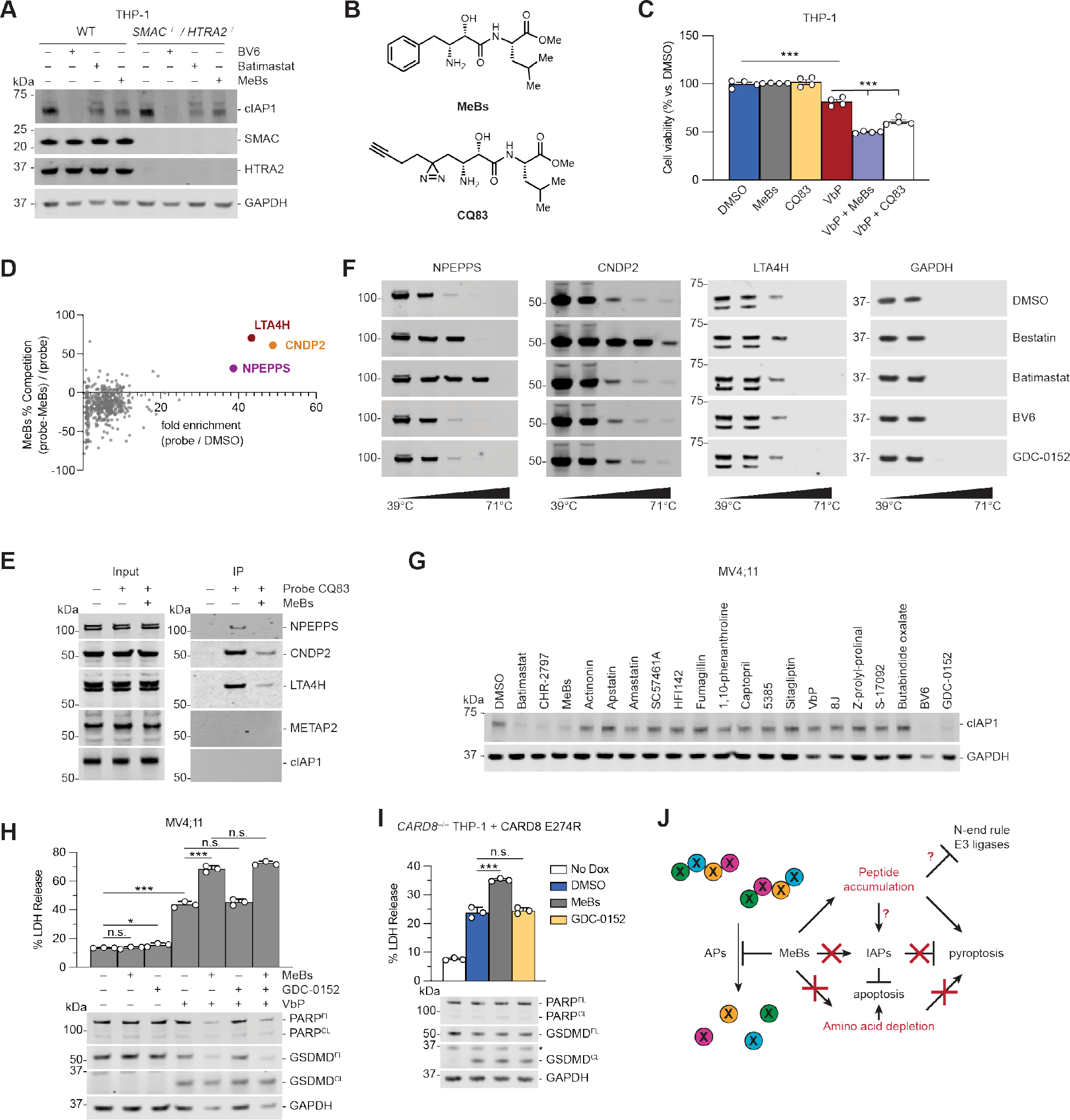
AP inhibition induces cIAP1 and NT degradation independently. (**A**) WT or *SMAC^−/−^/HTRA2^−/−^* THP-1 cells were treated with MeBs (10 µM), Batimastat (10 µM) or BV6 (5 µM) for 24h prior to immunoblot analysis. (**B**) Chemical structures of MeBs (top) and CQ83 (bottom). (**C**) THP-1 cells were treated with MeBs (1 µM), CQ83 (1 µM), and/or VbP (10 µM), and cell viability was assessed by CellTiter-Glo (CTG) after 6 h. (**D,E**) Scatter plot (**D**) and immunoblots (**E**) depict the proteins enriched by CQ83 and competed by MeBs as determined by TMT-MS (**D**) or immunoblotting (**E**). (**F**) CETSA analyses of bestatin (10 µM), Batimastat (10 µM), BV6 (5 µM) and GDC-0152 (5 µM) in HEK 293T cell lysates. (**G**) MV4;11 cells were treated with the indicated aminopeptidase inhibitors or IAP agonists (BV6, GDC-0152) for 24 h prior to immunoblotting analysis. All compounds were treated at 10 µM except: fumagillin (3 µM); compound 5385 (DPP7 inhibitor; 20 µM), compound 8j (selective DPP8/9 inhibitor; 20 µM), and sitagliptin (DPP4 inhibitor; 20 µM), VbP (2 µM), BV6 (5 µM), and GDC-0152 (5 µM). (**H**) MV4;11 cells were treated with VbP (10 µM), MeBs (10 µM), and/or GDC-0152 (5 µM) for 6 h prior to LDH and immunoblot analyses. (**I**) *CARD8*^−/−^ THP-1 cells stably containing doxycycline (DOX)- inducible CARD8 E274R were induced with 100 ng/mL DOX for 16 h prior to treatment with DMSO, MeBs (10 µM) or GDC-0152 (5 µM) for 6 h. Samples were then collected for LDH release and immunoblot analyses. (**J**) The proposed biological impacts of MeBs. MeBs inhibits APs, resulting in blockade of amino acid recycling and peptide accumulation. Some peptides bind to and thereby degrade cIAP1, some inhibits the N-end rule pathway, and some trigger CARD8/NLRP1 NT degradation. The peptides that modulate each of these effects are likely (at least partially) distinct. Data are means ± SEM of 3 or more replicates. All data, including immunoblots, are representative of three or more independent experiments. *** *p* < 0.001, * *p* < 0.05 by two-sided Students *t*-test. n.s., not significant. See also Figure S5 and Table S4.

To unbiasedly identify the direct protein targets of MeBs, and in particular to evaluate if MeBs itself associates with cIAP1, we synthesized a MeBs analog that contains a photoactivatable diazirine group and a click chemistry compatible alkyne handle to enable covalent attachment to and enrichment of target proteins (CQ83, **Fig. 6B**). Importantly, CQ83, like MeBs, enhanced VbP-induced pyroptosis in THP-1 cells (**Fig. 6C**), demonstrating that CQ83 still inhibits the relevant protein targets in cells. We next incubated HEK 293T lysates with CQ83 with or without MeBs, crosslinked CQ83 to target proteins using UV light, coupled CQ83-modified proteins to biotin using click chemistry, and enriched biotinylated proteins on streptavidin beads. The enriched proteins were then subjected to on-bead trypsinization, tandem mass tag (TMT)- labeling, and quantitative protein mass spectrometry analysis (**Fig. S5B**). As expected, we found that CQ83 selectively enriched several APs, including LTA4H, NPEPPS and CNDP2 (**Fig. 6D, Table S4**), and we confirmed these results by immunoblotting (**Fig. 6E**). In contrast, CQ83 did not enrich cIAP1 (**Fig. 6D,E**). Consistent with these data, bestatin (and, with differing selectivity, batimastat), but not BV6 and GDC-0152, stabilized these APs in a cellular thermal shift assay (CETSA) in HEK 293T cell lysates (**Fig. 6F**). Overall, these data indicate that AP inhibitors do not directly interact with cIAP1.

Instead, we hypothesized that AP inhibition indirectly caused cIAP1 degradation, likely by stabilizing peptides that interact with the BIR domains. We next tested a diverse panel of peptidase inhibitors to determine if any others similarly triggered cIAP1 degradation, and we discovered that only MeBs, CHR-2797, and batimastat induced this effect in MV4;11 cells (**Fig. 6G, Fig. S1D**). The SMAC mimetics BV6 and GDC-0152 were used as positive controls in this experiment. Notably, only these three peptidase inhibitors synergized with VbP to induce more pyroptosis (**Fig. S5C,D**) (Chui et al., 2019). Thus, inhibition of the same (or similar) APs accelerates the degradation of cIAP1 and CARD8/NLRP1. However, it remained unclear if cIAP1 degradation was involved in inflammasome activation, or if it was part of a distinct pathway downstream of AP inhibition.

To determine if cIAP1 degradation is involved in the induction of synergistic pyroptosis, we next evaluated the impact of GDC-0152 on pyroptosis in MV4;11 cells. GDC-0152 did not induce any GSDMD cleavage on its own, nor did it increase the amount of VbP- or VbP plus MeBs-induced GSDMD cleavage (**Fig. 6H**). In addition, GDC-0152, unlike MeBs, did not induce any additional cell death in *CARD8^−/−^* THP-1 cells ectopically expressing CARD8 E274R (**Fig. 6I**). Thus, cIAP1 degradation does not augment CARD8 inflammasome activation (**Fig. 6J**). Collectively, these data indicate that AP inhibition induces the accumulation of many cytosolic peptides, which in turn causes a myriad of biological responses, including N-end rule pathway blockade, cIAP1 degradation, and CARD8/NLRP1 NT degradation (**Fig. 6J**). In particular, we hypothesize that the accumulation of peptides with certain destabilizing N-terminal residues block the N-end rule E3 ligases, those with N-terminal IBM sequences induce the degradation of cIAP1, and those with distinct (but as yet unknown) sequences accelerate the degradation of NLRP1 and CARD8. Regardless, the MeBs-induced cIAP1 degradation pathway revealed here provides more evidence that peptide accumulation triggers several biological responses, even though this pathway is not involved in regulating NLRP1 and CARD8.

### Other proteotoxic stress inducers trigger synergistic pyroptosis

To more unbiasedly assess the cellular response to MeBs, we next performed RNA sequencing (RNA-seq) analysis on DMSO- and MeBs-treated *CASP1*^−/−^ MV4;11 cells (**Table S5**). We found that MeBs triggered the upregulation of several genes involved in the response to damaged or misfolded protein accumulation, or proteotoxic stress, including *ATF4*, *ATF5*, *TXNIP*, *TRIB3*, *DDIT4*, *ADM2*, *CHAC1*, *SESN2* (**Fig. 7A**) (Kovaleva et al., 2016; Mungrue et al., 2009; Pakos-Zebrucka et al., 2016; Yang et al., 2021). We confirmed the upregulation of several of these transcripts using quantitative PCR (qPCR) (**Fig. 7B**). It is well-established that peptides interfere with chaperone-mediated protein folding (Li et al., 2003; Otvos et al., 2000). Thus, we reasoned that accumulation of proteasome-generated peptides, and especially those with hydrophobic sequences, interfere with protein folding and thereby induce genes involved in proteotoxic stress. In addition to these genes, we found that MeBs upregulated the transcription of several genes encoding amino acid transporters, including *SLC38A2*, *SLC7A5*, and *SLC3A2* (**Fig. 7A, Table S5**). These results suggest that MeBs does cause some amino acid starvation, even it does not appreciably impact the overall amino acid levels (**Fig. 4C**). Notably, a previous study found that CHR-2797 induced a very similar transcriptional changes (Krige et al., 2008), indicating that these responses are indeed due to AP inhibition.

**Figure 7.**
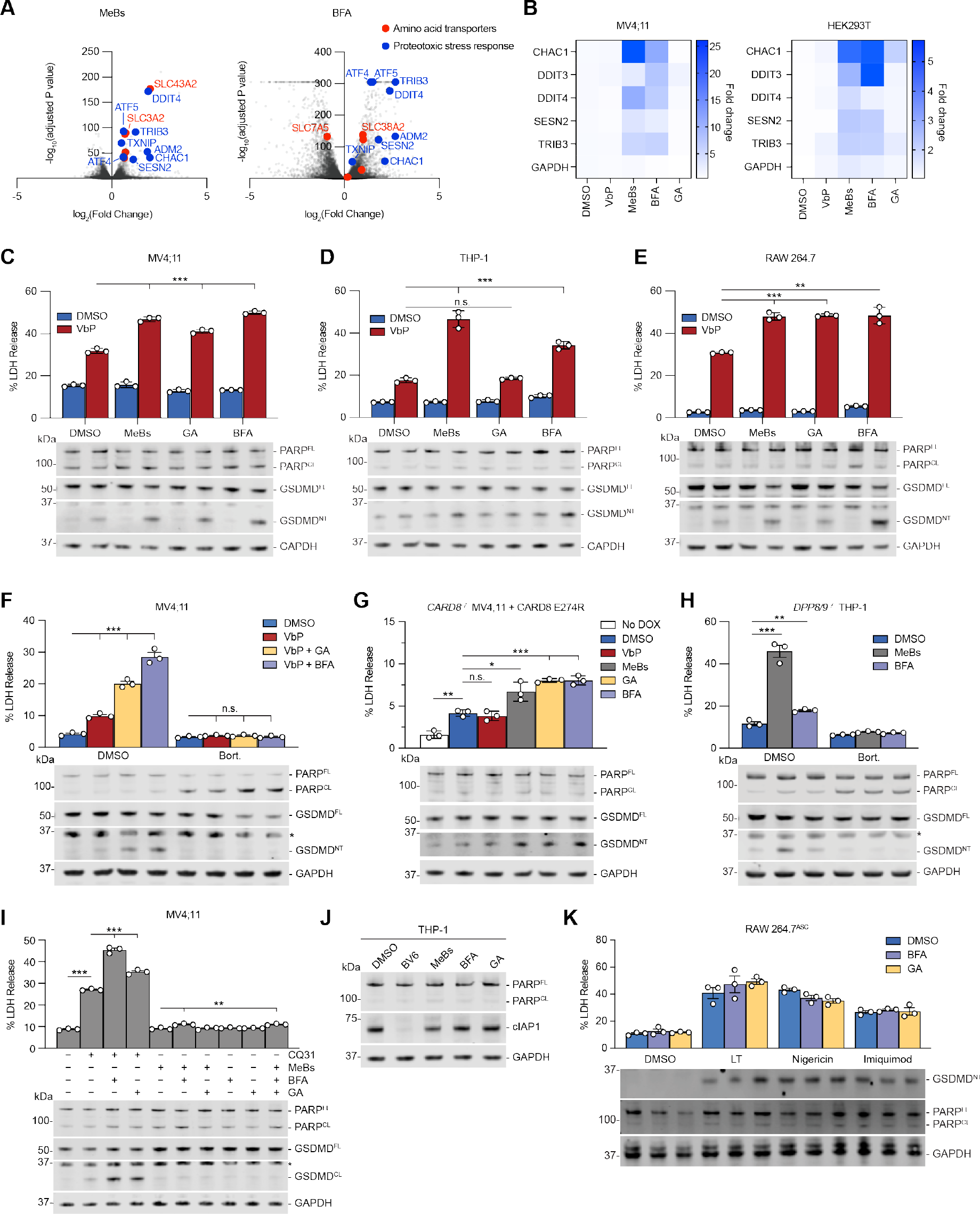
Proteotoxic agents accelerate NT degradation. (**A,B**) *CASP1^−/−^* MV4;11 and/or HEK 293T cells were treated with VbP (10 µM), MeBs (10 µM), brefeldin A (BFA; 1.78 µM) or Geldanamycin (GA; 1 µM) for 6 h before gene expression relative to a DMSO control was determined by RNA-seq (**A**) or RT-qPCR (**B**). (**C**-**F**) The indicated cell lines cell lines were treated with MeBs (10 µM), GA (1 µM), BFA (1.78 µM), Bort. (10 µM) and/or VbP (10 µM) for 6 h prior to LDH and immunoblot analyses. (**G**) *CARD8^−/−^* MV4;11 cells stably expressing doxycycline (DOX)- inducible CARD8 E274R are treated with DOX (1 µg/mL), VbP (10 µM), MeBs (10 µM), GA (1 µM), or BFA (1.78 µM) for 6 h prior to LDH and immunoblot analyses. (**H**) *DPP8^−/−^/DPP9^−/−^* THP- 1 cells were treated with MeBs (10 µM), BFA (1.78 µM) and/or Bort. (10 µM) for 8 h prior to LDH release and immunoblot analyses. (**I**) MV4;11 cells were treated with CQ31 (20 µM) for 22 h, and/or MeBs (10 µM), GA (1 µM), or BFA (1.78 µM) for 6 h prior to LDH and immunoblot analyses. (**J**) THP-1 cells were treated with BV6 (5 µM) MeBs (10 µM), GA (1 µM), or BFA (1.78 µM) for 6 h prior to immunoblot analysis. Data are means ± SEM of 3 replicates. (**K**) RAW 264.7 cells stably expressing ASC were treated with LPS (5 µg/mL) for 16 h prior to treatment with BFA (1.78 µM) for 6 h, GA (1 µM) for 6 h, anthrax lethal factor and protective antigen (LT; 1 µg/mL each) for 6 h, nigericin (10 µM) for 3 h and/or imiquimod (100 µM) for 3 h. LDH and immunoblot analyses were performed. All data, including immunoblots, are representative of three or more independent experiments. *** *p* < 0.001, ** *p* < 0.01, * *p* < 0.05 by two-sided Students *t*-test. N.s., not significant. See also Figure S6 and Table S5.

We next wondered if other well-characterized agents that interfere with protein folding, including brefeldin A (BFA) and geldanamycin (GA), might similarly accelerate NLRP1 and CARD8 NT degradation. BFA inhibits protein export from the endoplasmic reticulum and thereby induces the unfolded protein response (UPR) (Citterio et al., 2008; Fujiwara et al., 1988; Helms and Rothman, 1992). GA inhibits the ATPase activity of heat-shock protein 90 (HSP90) and thereby destabilizes HSP90 client proteins (Neckers et al., 1999a; Neckers et al., 1999b). Notably, BFA upregulated the transcription of several of the same proteotoxic stress response genes as MeBs, including *ATF4*, *ATF5*, *TRIB4*, and *DDIT4* (**Fig. 7A**,**B, Table S5**), indicating at least some similarities between stresses caused these agents. However, GA did not appreciably alter mRNA transcript levels of these genes (**Fig. 7B**). Consistent with our hypothesis, we discovered that BFA, like MeBs, strongly synergized with VbP to induce more pyroptosis in MV4;11, THP-1, RAW 264.7, and OCI-AML2 cells (**Fig. 7C-E**, **Fig. S6A**), although it did not induce the release of significantly more IL-1β in N/TERT-1 keratinocytes (**Fig. S6B**). We also observed that GA had at least some synergy with VbP in MV4;11, RAW 264.7 cells, and N/TERT-1 keratinocytes, albeit not in THP-1 cells (**Fig. 7C-E, Fig. S6B**). It is not clear why BFA and GA do not synergize with VbP in N/TERT-1 keratinocytes and THP-1 cells, respectively, but we speculate it could be due to different proteostasis networks in the cell types.

As expected, both BFA- and GA-induced synergistic pyroptosis in MV4;11 was proteasome degradation-dependent, as bortezomib completely abolished this pyroptotic death (**Fig. 7F**). Moreover, BFA and GA induced more proteasome-dependent pyroptosis in *CARD8^−/−^* MV4;11 cells ectopically expressing the DPP9 non-binding CARD8 E274R mutant protein (**Fig. 7G, Fig. S6C**). Similarly, BFA induced more GSDMD cleavage in both *DPP8/9^−/−^* THP-1 cells and THP-1 cells expressing CARD8 E274R (**Fig. 7H, Fig. S6D**). In addition, both BFA and GA synergized with CQ31 to induce more pyroptosis in MV4;11 cells (**Fig. 7I**). Notably, we found that GA and BFA, unlike MeBs, did not induce cIAP1 degradation, consistent with their mechanisms not involving the stabilization of peptides (**Fig. 7J**). Despite these distinct mechanisms of action, however, it should be noted that the combination of MeBs, GA, and BFA did not induce pyroptosis without a DPP8/9-binding ligand (**Fig. 7I, S6E**). Overall, these data show that BFA and GA, like MeBs, accelerate NLRP1 and CARD8 NT degradation, but do not alone induce inflammasome activation because the DPP8/9 complexes sequester the freed CT fragments.

We next wanted to test the impact of BFA and GA on inflammasome activation by other stimuli. We found that BFA and GA did not impact LT-induced NLRP1B-dependent pyroptosis, imiquimod-induced NLRP3-dependent pyroptosis, nor nigericin-induced NLRP3-dependent pyroptosis in RAW 264.7 cells stably expressing ASC (RAW 264.7 cells do not endogenously express ASC, which is required for NLRP3 inflammasome formation) (**Fig. 7K**). These results confirm that these proteotoxic drugs do not non-specifically increase pyroptosis induced by all inflammasome triggers.

As translation is a major source of unfolded proteins, we next evaluated the impact of the translation inhibitor cycloheximide (CHX) on this pyroptotic pathway. As expected, CHX, but not VbP, MeBs, batimastat, GA, or BFA, greatly slowed the overall translation rate in HEK 293T cells, as measured by puromycin incorporation into nascent polypeptides (**Fig. S6F**.) Intriguingly, we found that CHX attenuated VbP and VbP plus MeBs-induced pyroptosis in MV4;11 cells (**Fig. S6G, H**), MeBs-induced pyroptosis in *PEPD^−/−^* THP-1 cells (**Fig. S2D**), as well as VbP-induced pyroptosis in primary rat and mouse macrophages (**Fig. S6I,J**). In addition, we found that CHX also rescued BFA- and GA-induced synergistic pyroptosis with VbP in MV4;11 cells (**Fig S6K**). We hypothesize that CHX reduces the amount of unfolded polypeptides in the cell, thereby reducing the overall burden for the chaperone machinery. However, translation blockade could obviously block pyroptosis via a variety of other mechanisms as well, for example by inhibiting the synthesis of proteins involved in the pyroptotic pathway. Nevertheless, these data collectively indicate that agents that interfere with protein folding, including peptides, accelerate the degradation of the NLRP1 and CARD8 NT fragments.

## DISCUSSION

The biological purposes of the NLRP1 and CARD8 inflammasomes have not yet been established (Bachovchin, 2021). Notably, NLRP1 is highly polymorphic, and an array of dissimilar pathogen-associated stimuli, including viral proteases and dsRNA, induce its activation in experimental systems (Hornung et al., 2009; Robinson et al., 2020; Tsu et al., 2020). Curiously, however, none of these stimuli universally activate all functional NLRP1 alleles in rodents and humans. Based on these findings, some have hypothesized that NLRP1 is rapidly evolving to detect distinct danger signals and does not have a single primordial function. Alternatively, we have proposed that both NLRP1 and CARD8 evolved to detect, albeit with different thresholds, a single unknown danger state that is linked in some way to DPP8/9 (**Fig. 1D**) (Bachovchin, 2021; Chui et al., 2020). Here, we now show NLRP1 and CARD8 detect protein misfolding that is associated with peptide accumulation (**Fig. S7A**).

NLRP1 and CARD8 sense the stability of their NT fragments to proteasome-mediated degradation, but the cellular factors that regulate this stability are poorly understood. In this Article, we demonstrate that several distinct and well-characterized small molecules, including MeBs, BFA, and GA, accelerate the rate of NT degradation. Notably, MeBs, BFA, and GA stabilize oligopeptides, block protein export, and interfere with chaperone function, respectively, and therefore act via different molecular mechanisms. Nevertheless, all three agents interfere with protein folding, thereby destabilizing the NT fragments (**Fig. S7A**). It should be noted that, even though these proteotoxic drugs accelerate NT degradation, they do not cause inflammasome activation on their own because the DPP8/9 ternary complex quenches the released CT fragments; a DPP8/9-binding ligand (e.g., VbP or an XP-containing peptide) is needed to destabilize the ternary complex and enable inflammasome assembly to proceed. We hypothesize that imposing even greater proteotoxic stress, for example by combining these drugs with an agent that impacts protein folding in a different way, might trigger inflammasome activation without having to simultaneously add an exogenous DPP8/9 inhibitor. On that note, we recently discovered that reductive stress appears to contribute to NLRP1 inflammasome activation (Ball et al., 2021), and we speculate that certain antioxidants might interfere with protein folding by such a distinct mechanism (Tu and Weissman, 2004; Wang et al., 2022).

We should emphasize that our results here, coupled with our previous work (Rao et al., 2022), show that peptides play important roles in controlling NLRP1 and CARD8 activation both upstream and downstream of the proteasome (**Fig. 1D**, **Fig. S7A**). As described in this manuscript, a large and diverse array of (likely hydrophobic) peptides, which are stabilized by MeBs (and probably to lesser extent by VbP and CQ31), accelerate the rate of NT degradation (**Fig. S7A**). Peptides with N-terminal XP sequences, which are stabilized by CQ31 and VbP but not MeBs, disrupt the DPP8/9-containing repressive ternary complex. Moreover, VbP itself essentially mimics an extremely high concentration of XP peptides. Thus, we propose that the overall intracellular peptide pool increases NT degradation, and that DPP8/9 sense XP peptide levels as a checkpoint to verify that NT degradation is indeed associated with peptide build-up. A possible reason that XP-containing peptides are monitored is discussed below. It should be noted that XP peptides (i.e., CQ31-stabilized peptides) alone are sufficient to activate CARD8, but that a large set of peptides including XP-containing peptides (i.e., both MeBs- and CQ31-stabilized peptides) are required to activate NLRP1 (**Fig. 2I**). These data are consistent with the idea that NLRP1 is more inflammatory than CARD8 and therefore has a higher threshold for activation (Bachovchin, 2021; Ball et al., 2020).

Projecting forward, at least two questions remain unanswered: 1) Why does the impairment of protein folding accelerate the degradation of the NT fragments, and 2) why are XP peptides so important? Regarding the first question, we hypothesize that unfolded polypeptides interfere with the initial folding of nascent NLRP1/CARD8 protein and/or unravel already folded NLRP1/CARD8. As the proteasome rapidly destroys misfolded proteins (Baugh et al., 2009; Liu et al., 2003), this would lead to their rapid degradation. If correct, it seems likely the polymorphic and disordered NT regions exist to modulate the propensity of the ZU5 domains to misfold (Chui et al., 2020; Waldo et al., 1999). Regarding the second question, we speculate that XP- containing peptides might serve as an early warning sign of impending proteostasis failure. Briefly, defective protein folding will cause many newly misfolded proteins to be degraded by the proteasome, thereby generating peptides. If these peptides are so abundant that they overwhelm the ability of APs to destroy them, they may then initiate a disastrous feedforward cycle, in which the peptides in turn interfere with protein folding, generate more peptides, and so on, ultimately leading to proteostasis failure (**Fig. S7B**). XP-containing peptides are especially challenging to cleave due to their conformationally restricted proline bond, and thus might accumulate first and serve as a harbinger of catastrophe.

Finally, we should note that NLRP1 (and possibly the CARD8) did not evolve to sense small molecules that interfere with proteostasis, but rather the presence of infectious pathogens. Nevertheless, this work showcases the utility of fast acting and selective chemical probes for deconvoluting complex immunological pathways. We anticipate that the development of additional chemical tools, likely including ones that induce reductive stress (Wang et al., 2022), will enable further delineation of molecular mechanisms that regulate NLRP1 and CARD8 activation and ultimately help illuminate their relationship with infectious agents. On that note, the polymorphic rodent NLRP1 alleles appear to have similar relative sensitivities to *Toxoplasma gondii* (*T. gondii*) infection and VbP (Cirelli et al., 2014; Ewald et al., 2014; Gai et al., 2019), and we therefore speculate that *T. gondii* may compromise protein folding through as yet unknown mechanisms. We expect that future studies with *T. gondii* and potentially other pathogens, coupled with the use of high-quality chemical probes, will reveal the full biological purpose of these enigmatic inflammasomes.

## Acknowledgements

This work was supported by the NIH (R01 AI137168, R01 AI163170, and R01 CA266478 to D.A.B.; T32 GM007739-Andersen to A.R.G; NIH T32 GM115327-Tan to E.L.O.-H.; F30 CA008748 to A.R.G.; the MSKCC Core Grant P30 CA008748), Gabrielle’s Angel Foundation (D.A.B.), Mr. William H and Mrs. Alice Goodwin, the Commonwealth Foundation for Cancer Research, and The Center for Experimental Therapeutics of Memorial Sloan Kettering Cancer Center (D.A.B.), the Emerson Collective (D.A.B.), and the Marie-Josée Kravitz Women in Science Endeavor (WISE) fellowship (SDR).

## Author Contributions

D.A.B. conceived and directed the project. E.L.O.-H., H.-C.H., S.D.R., Q.W., Q.C., C.M.O., A.J.C., A.R.G., A.B., and D.P.B. performed cloning, gene editing, biochemistry, and cell biology experiments. Q.C. synthesized CQ83 and performed chemoproteomics experiments and validation. E.L.O.-H., H.-C.H., J.R.C., and M.S. performed and analyzed metabolomics experiments. D.A.B., E.L.O.-H., and H.-C.H. wrote the manuscript.

## Materials and Methods

### METHOD DETAILS

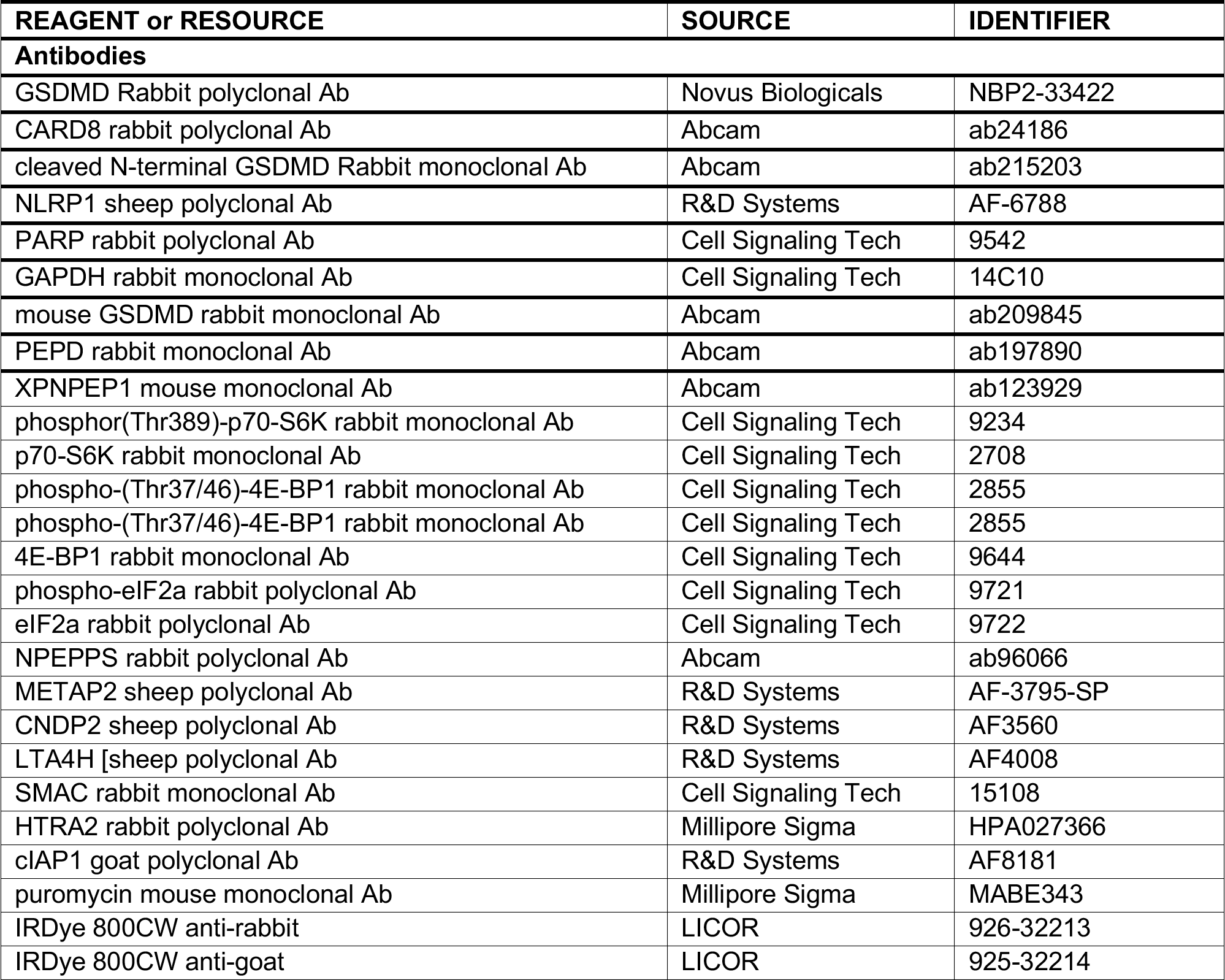
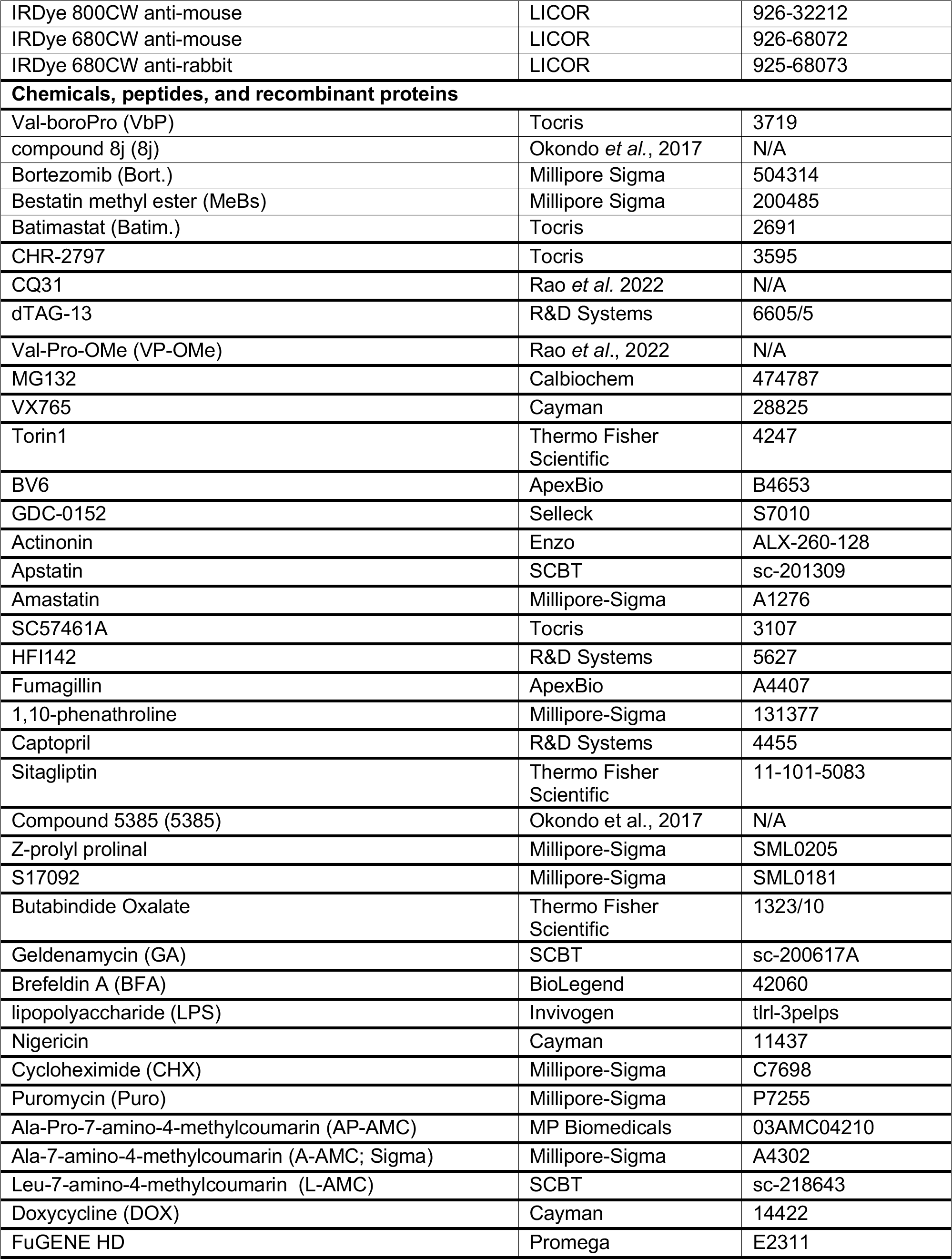
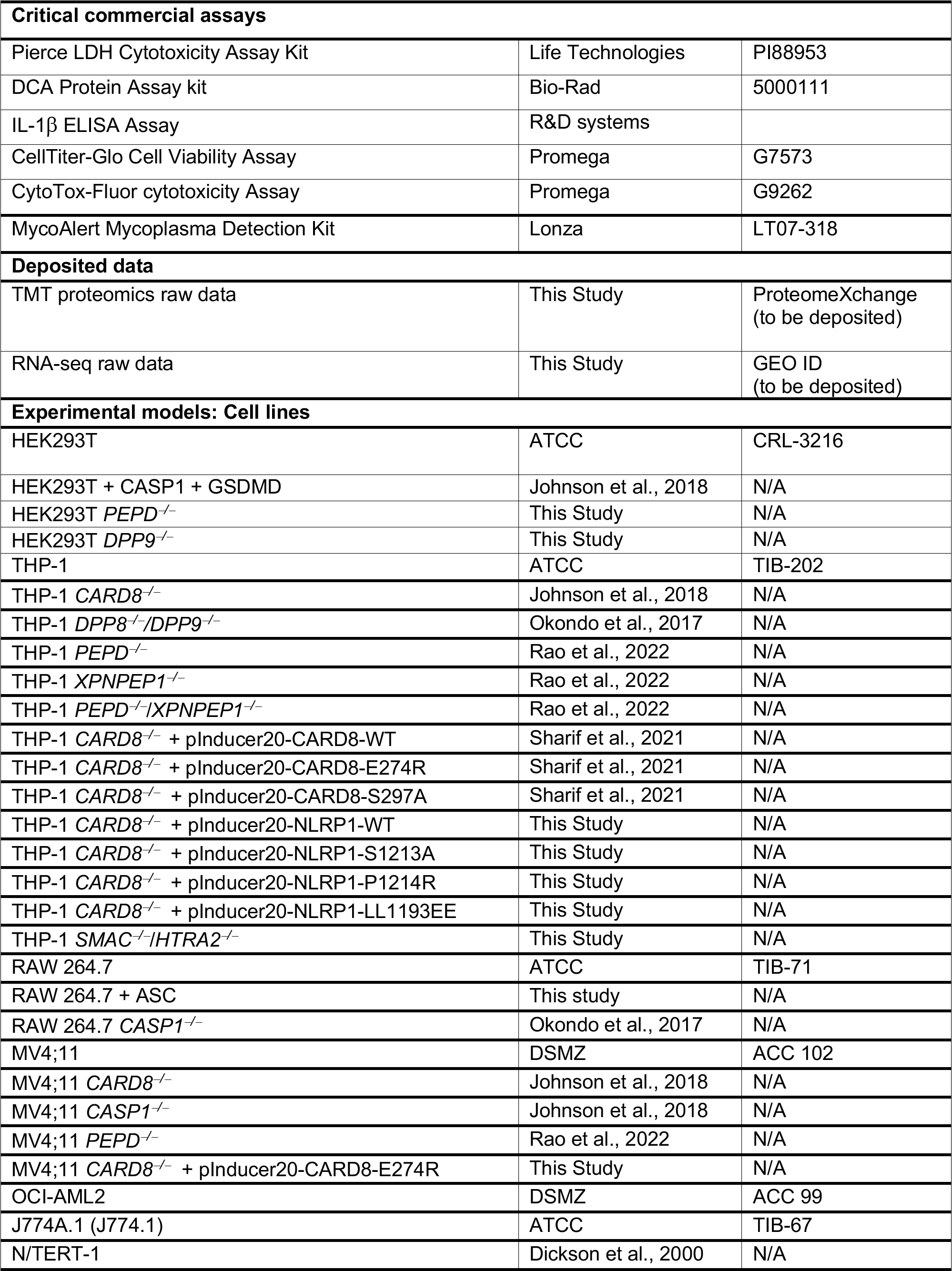
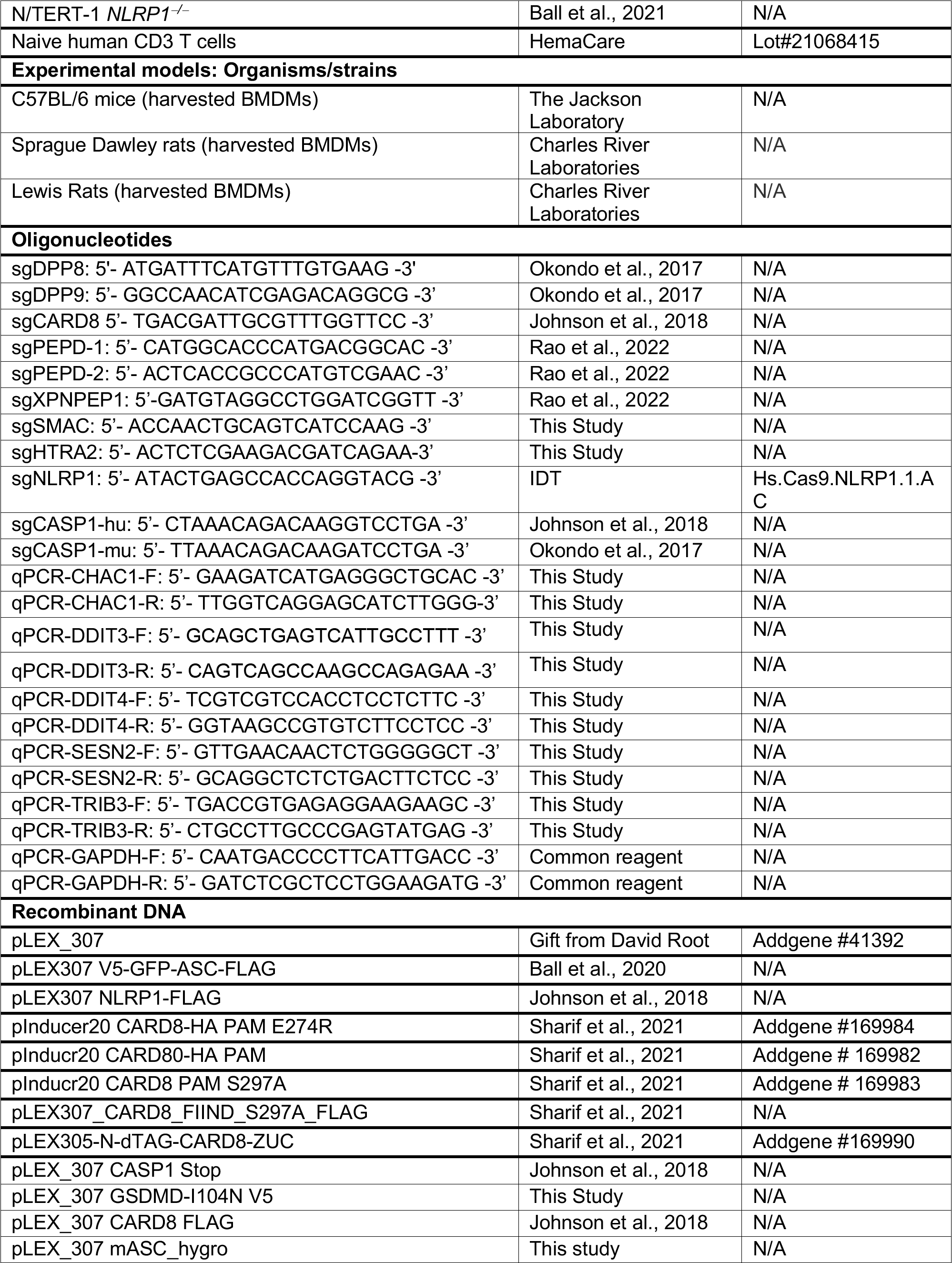
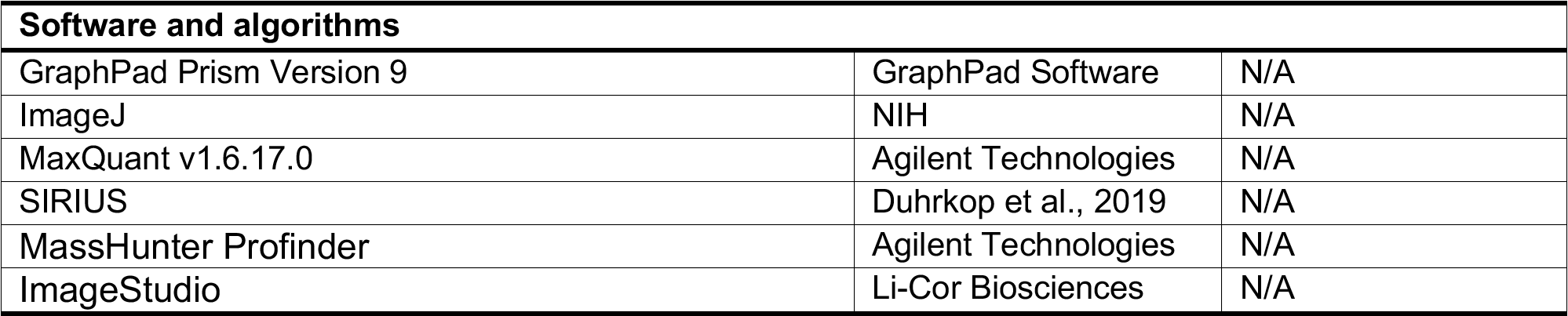

#### Cell Culture

HEK 293T, THP-1, J774.1 and RAW 264.7 cells were purchased from ATCC. OCI-AML2 and MV4;11 cells were purchased from DSMZ. Naïve CD3 human T cells were purchased from HemaCare (Lot #21068415). N/TERT-1 cells were a gift from the Rheinwald Lab (Dickson et al., 2000). HEK 293T, RAW 264.7 and J774.1 cells were grown in Dulbecco’s Modified Eagle’s Medium (DMEM) with L-glutamine and 10% fetal bovine serum (FBS). Naïve human CD3 T cells, THP-1, MV4;11, and OCI-AML2 cells were grown in Roswell Park Memorial Institute (RPMI) medium 1640 with L-glutamine and 10% FBS. N/TERT-1 cells were grown in Keratinocyte serum free medium (KSFM) supplemented with 1X penicillin/streptomycin, bovine pituitary extract (25 µg/ml) and epidermal growth factor (EGF) (0.2 ng/mL). All cells were grown at 37°C in a 5% CO2 atmosphere incubator. Cell lines were regularly tested for mycoplasma using the MycoAlert Mycoplasma Detection Kit (Lonza). *CARD8^−/−^*, *DPP8^−/−^*/*DPP9^−/−^*, *PEPD^−/−^*, *PEPD^−/−^*/*XPNPEP1^−/−^*, and *XPNPEP1^−/−^* THP-1 cells, *CASP1^−/−^* RAW 264.7 cells, *CARD8^−/−^*, *PEPD^−/−^*, and *CASP1^−/−^* MV4;11 and *NLRP1^−/−^* N/TERT-1 cells were generated as previously described (Ball et al., 2021; Johnson et al., 2018; Okondo et al., 2017). Doxycycline (DOX)-inducible CARD8 and NLRP1 WT and mutant knock-in *CARD8^−/−^* THP-1 were generated as previously described (Hollingsworth et al., 2021; Sharif et al., 2021), and DOX-inducible CARD8 knock-ins to *CARD8^−/−^* MV4;11 cells were similarly generated for this study. Briefly, *CARD8^−/−^* MV4;11 cells were infected with the indicated lentivirus containing pInducer20 construct, followed by selection with G418 (Geneticin) at 400 μg/mL until all control cells were dead (approximately 14 days).

#### Cloning

Plasmids for CARD8 WT and variants, CASP1, GSDMD, NLRP1 WT and variants, ASC-GFP, dTAG-CARD8^ZUC^ were cloned as described previously (Hollingsworth et al., 2021; Johnson et al., 2018; Sharif et al., 2021). Briefly, DNA sequences encoding the genes were purchased from GenScript, amplified by polymerase chain reaction (PCR), shuttled into the Gateway cloning system (ThermoFisher Scientific) using pDONR221 and pLEX307 vectors originating from pLEX307 (Addgene #41392). sgRNAs were designed using the Broad Institute’s web portal (Doench et al., 2016) (http://www.broadinstitute.org/rnai/public/analysis-tools/sgrna-design) and cloned into the lentiGuide-puro vector (Addgene #52963) as described previously (Sanjana et al., 2014). The sgRNA sequences used are described in the STAR Methods table.

#### CellTiter-Glo cell viability and CytoTox-Fluor cell death assays

Cells were plated (2,000 cells per well) in white, 384-well clear-bottom plates (Corning) using an EL406 Microplate Washer/Dispenser (BioTek) in 25 μL final volume of medium. To the cell plates were added compounds at different concentrations using a pin tool (CyBio) and the plates were allowed to incubate for 1 h in the incubator before adding VbP (10 µM). After incubation for indicated times, CytoTox-Fluor reagent (Promega, G9262) was added according to the manufacturer’s protocol. The assay plates were then incubated for another 30 min before fluorescence was recorded using a Cytation 5 Cell Imaging Multi-Mode Reader (BioTek). Next, CellTiter-Glo reagent (Promega, G7573) was subsequently added to the assay plates following the manufacturer’s protocol. Assay plates were shaken on an orbital shaker for 2 min and incubated at 25 °C for 10 min. Luminescence was then read using a Cytation 5 Cell Imaging Multi-Mode Reader (BioTek).

#### LDH cytotoxicity assays

HEK 293T cells were transiently transfected and treated with inhibitors as indicated. MV4;11, THP-1, OCI-AML2, RAW 264.7 or J774.1 cells were plated in 12-well culture plates at 5 × 10^5^ cells/well, Naïve human CD3 T cells were plated in 12-well tissue culture plates at 2 × 10^6^ cells/well, and N/TERT-1 cells were plated at 2 × 10^5^ cells/well and treated with chemical compounds as indicated. Supernatants were analyzed for LDH activity using the Pierce LDH Cytotoxicity Assay Kit (Life Technologies). LDH activity was quantified relative to a lysis control where cells were lysed by adding 8 µL of a 9% Triton X-100 solution.

#### Immunoblotting

Cells were washed 2 × in PBS (pH = 7.4), resuspended in PBS, and lysed by sonication. Protein concentrations were determined and normalized using the DCA Protein Assay kit (Bio-Rad). Samples were run on NuPAGE 4 to 12%, Bis-Tris, 1.0 mm, Midi Protein Gel (Invitrogen) for 45–60 min at 175 V. Gels were transferred to nitrocellulose with the Trans-Blot Turbo Transfer System (Bio-Rad). Membranes were blocked with Intercept (TBS) Blocking Buffer (LI-COR) for 30 min at ambient temperature, before incubating with primary antibody overnight at 4 °C. Blots were washed 3 times with TBST buffer before incubating with secondary antibody for 60 min at ambient temperature. Blots were washed 3 times, rinsed with water and imaged via Odyssey CLx (LI-COR). Antibodies are listed in the Key Resources Table.

#### CRISPR/Cas9 gene editing

*DPP8/9*, *CARD8*, *PEPD*, *XPNPEP1*, and *PEPD/XPNPEP1* knockout THP-1 cell lines and all HEK 293T, RAW 264.7, N/TERT-1 and MV4;11 knockout cell lines were generated as previously described (Ball et al., 2021; Johnson et al., 2018; Okondo et al., 2017; Rao et al., 2022; Sharif et al., 2021). Briefly, 5 x 10^5^ HEK 293T cells stably expressing Cas9 were seeded in 6-well tissue culture dishes in 2 mL of media per well. The next day cells were transfected according to manufacturer’s instructions (FuGENE HD, Promega) with 2 μg of the sgRNA plasmid(s). After 48 h, cells were transferred to a 10 cm tissue culture dish and selected with puromycin (1 μg/mL) until control cells were all dead. Single cell clones were isolated by serial dilution and confirmed by Western blot or sequencing, as indicated. To generate knockouts in RAW 264.7, MV4;11 and THP-1 cells, 1.5 x 10^6^ cells stably expressing Cas9 (Johnson et al., 2018) were infected with lentivirus containing sgRNA plasmids. After 48 h, cells were selected with puromycin (1 μg/mL) or hygromycin (100 μg/mL). Single cell clones were isolated by serial dilution and confirmed by Western blotting. *NLRP1* knockout N/TERT-1 keratinocytes were prepared by using the Neon Transfection System (ThermoFisher Scientific) following the manufacturer’s recommendations to deliver Cas9 ribonucleoprotein complexes containing an Alt-R CRISPR-Cas9 sgRNA and recombinant Cas9 (IDT). Briefly, sgRNA complexes were prepared by combining predesigned Alt-R CRISPR-Cas9 crRNA (NLRP1: 5’- ATACTGAGCCACCAGGTACG -3’) with Alt-R CRISPR-Cas9 tracrRNA to 44 µM and annealing by heating to 95 °C for 5 min followed by gradual cooling to ambient temperature over 30 min. To form the RNP complexes sgRNA samples and recombinant SAlt-R Cas9 enzyme were combined and incubated for 20 min.

#### Stable Cell Line Generation

Cells stably expressing indicated protein constructs were generated by infection with lentivirus containing the desired plasmids. Briefly, the lentivirus was produced by transfecting 70% confluent HEK 293T cells with the desired plasmid along with psPAX2 and pMD2.G following the manufacturer’s instructions (FuGENE HD, Promega). The virus-containing medium was collected 48 h after transfection, passed through a 0.45 µm filter, and concentrated by PEG precipitation (Abcam). THP-1, RAW 264.7, or MV4;11 cells were infected with the prepared lentivirus by centrifuging at 1000 x g for 1 h. After 48 h of incubation, cells were selected with an appropriate antibiotic.

#### Transient transfections

HEK 293T cells were plated in 6-well culture plates at 5.0 × 10^5^ cells/well in DMEM. The next day, the indicated plasmids were mixed with an empty vector to a total of 2.0 µg DNA in 125 µL Opti-MEM and transfected using FuGENE HD (Promega) according to the manufacturer’s protocol. Unless indicated otherwise, 0.05 µg *CASP1*, 0.025 µg *CARD8* and 0.025 µg *GSDMD*-I104N plasmids were used. After 16-20 h, the cells were treated as described. For microscopy experiments, HEK 293T cells were seeded into Lab-Tek II 8-well chambered coverglass plates at 2 × 10^4^ cells per chamber. After 48 h, the cells were transfected with 0.02 µg of plasmids encoding C-terminally FLAG-tagged *NLRP1*, 0.01 µg of a plasmid encoding N-terminally V5-GFP-tagged *ASC*, and 0.37 µg of a plasmid encoding *RFP* using FuGene as the transfection agent and given 24 h to express protein and then treated with the indicated agent for 24 h, Hoechst stain (1 µg/mL) was added.

#### Fluorescence microscopy and analysis

Cells transfected as described above were imaged on a Zeiss Axio Observer.Z1 inverted widefield microscope using ×10/0.95NA air objective. For each well, 10 positions were imaged in the brightfield, Hoechst (DAPI), RFP, and GFP channels. Data was analyzed using custom macro written in ImageJ/FIJI. The number of cells containing GFP-ASC specks was quantified by setting threshold values on the GFP channel, and performing the ‘Analyze Particles’ algorithm, size = 0–∞ and circularity = 0.50–1.00. The data was then exported to spreadsheet software, analyzed to compute the ratio of specks.

#### Substrate assays

For the peptide assay, a solution of substrate (PASKYLF) was synthesized by standard solid state peptide synthesis methods using chlorotrityl resin, and prepared in water. Lysates (normalized to 0.5 mg/mL by DC Assay kit (Bio-Rad) were added to a 384-well, black, clear-bottom plate (Corning) with peptide PASKYLF (final conc. 1 mM), with a final volume per well of 25 μL. Alanine liberated was measured as increasing fluorescence signal (Resorufin, Ex/Em: 535/587 nm) recorded at 25 °C using an L-alanine assay kit (Abcam, ab83394) at 25 °C according to manufacturer’s instructions. For the AMC reporter assays, experiments were performed in cells. For in cell assays, 1.0 x 10^5^ *CARD8*^−/−^ THP-1 or HEK 293T WT cells were seeded per well in a 96-well, black, clear-bottom plate (Corning) in Opti-MEM reduced serum media and treated with compounds for 6 h before substrate (Ala-AMC, 100 μM; Leu-AMC, 100 μM; Ala-Pro-AMC, 250 μM) was added to the media to initiate the reaction. Substrate cleavage was measured as increasing fluorescence signal (Ex/Em: 380/460 nm) recorded at 25 °C for 25- 40 mins. Cleavage rates are reported as the slope of the linear regression of AMC fluorescence vs. time data curve.

#### CETSA analysis

HEK 293T cells were homogenized by sonication and cleared of debris by centrifugation at 10,000 x g for 10 min. Clarified lysates were then incubated with indicated inhibitors for 30 min, before heating at indicated temperatures for 30 min. Aggregated proteins were then precipitated by centrifugation at 18,000 x g for 20 min. Supernatant was then harvested and then subjected to immunoblotting.

#### Metabolite analysis using LC-MS

HEK 293T (0.5 x 10^6^) or THP-1 (0.75 x 10^6^) cells were seeded on 6-well tissue culture dishes in 2 mL of DMEM or RPMI, respectively, supplemented with 10% FBS per well. The next day, cells were treated with the indicated compounds for up to 6 h. Metabolism was quenched and metabolites were extracted by aspirating medium and adding 1 mL of ice-cold 80:20 methanol:water containing ^13^C amino acid standards-. After overnight incubation at -80 °C, cells were collected and centrifuged at 20,000 x g for 20 min at 4 °C. The supernatants were dried in a vacuum evaporator (Genevac EZ-2 Elite) for 3 hours. Dried extracts were resuspended in 40 µL of 60% acetonitrile in water. Samples were vortexed, incubated on ice for 20 min, followed by addition of 10 µL of methanol. After briefly vortexing, the dried extracts were clarified by centrifugation at 20,000 x g for 20 min at 4°C.

Dipeptide analysis was achieved using an Agilent 6545 Q-TOF mass spectrometer with Dual Jet Stream source in positive ionization, coupled to an Acquity UPLC BEH Amide column (150 mm x 2.1 mm, 1.7 µm particle size, Waters) kept at 40 °C. Composition of Mobile Phase A consisted of 10 mM ammonium acetate in 10:90 acetonitrile: water with 0.2% acetic acid at pH 4. Mobile Phase B consisted of 10 mM ammonium acetate in 90:10 acetonitrile: water with 0.2% acetic acid at pH 4. The gradient was as follows at an initial flow rate of 0.4 mL/min at 95% B; 9 min, 70% B; 13 min, 30% B; 14 min, 30% B; 14.5 min, 95% B. The flow rate was increased to 0.6 mL/min from 15 min, 95% B to 20 min, 95% B. The injection volume was 5 µL for each sample. MS parameters included: gas temp: 300 °C; gas flow: 10 L/min; nebulizer pressure: 35 psig; sheath gas temp: 350 °C, sheath gas flow: 12 L/min; VCap: 4000 V; fragmentor: 125 V. Data was acquired from 50 – 1700 m/z with reference mass correction (m/z: 121.05087 and 922.00980). Identification of dipeptides was achieved by running synthesized dipeptide standards to confirm retention time matching as well as performing tandem mass spectrometry (MS/MS) for spectral matching to SIRIUS (Duhrkop et al., 2019). Data analysis was performed using Agilent MassHunter Profinder 10.0 (Agilent Technologies).

Measurement of amino acid levels in *PEPD^−/−^*, *DPP8/9^−/−^* and WT HEK 293T cells was achieved by plating 6.5 x 10^5^ cells and seeding overnight, followed by treatment with VbP or MeBs for 3 h, then washing and harvesting of cells by pipetting in phosphate buffered saline, of which a small aliquot was collected for cellular protein normalization. Metabolite analysis was performed by Metabolon.

#### Stable Isotope Tracing

HEK 293T (0.5 x 10^6^) were grown for 23 days in labeled media consisting of DMEM deficient in L-leucine and L-glutamine, then supplemented with 10% dialyzed fetal bovine serum (FBS) and ^13^C6-leucine (^13^C-Leu) and ^13^C5-glutamine (^13^C-Gln). 0.5 x 10^6^ cells were plated in labeled media, then the following day labeled media was replaced with unlabeled media (DMEM supplemented with 10% FBS) and simultaneously treated with VbP (10 µM), MeBs (10 µM) or Bortezomib (10 µM) for 6 h. Metabolism was quenched and metabolites were extracted by aspirating medium and adding 1 mL of ice-cold 80:20 methanol:water. After overnight incubation at -80 °C cells were collected and centrifuged at 20,000 x g for 20 min at 4 °C. The supernatants were dried in a vacuum evaporator (Genevac EZ-2 Elite) for 3 hours. Dried extracts were resuspended in 40 µL of 60% acetonitrile in water.

Samples were vortexed, incubated on ice for 20 min, followed by addition of 10 µL methanol. After briefly vortexing, the dried extracts were clarified by centrifugation at 20,000 x g for 20 min at 4 °C.

Amino acid isotope tracing analysis was achieved using an Agilent 6545 Q-TOF mass spectrometer with Dual Jet Stream source in positive ionization using the same liquid chromatography gradient and mass spectrometry conditions previously mentioned. Data analysis was performed using Agilent MassHunter Profinder 10.0 for isotope tracing data where natural isotope abundance correction was performed (Agilent Technologies

#### Diazirine crosslinking and click chemistry

HEK 293T cells were harvested and pelleted at 400 x g, washed 3 x with cold PBS, resuspended in 1 mL of PBS and lysed by sonication. Lysates were then clarified at 20,000 x g for 15 min. The soluble fraction was retained, and protein concentrations were determined using the DC Protein Assay kit (Bio-Rad) and adjusted to 1 mg/mL. Aliquots of 1.0 mL were treated with CQ83 probe (10 µM) for 30 min. For competition experiments the lysates pretreated with 100 µM inhibitor MeBs (30 min) and then probe (30 min). Samples were then irradiated for 30 min in the Photochemical Reactor equipped with 350 nm lamps at 4 °C. 10 µL 4% SDS was added to each sample and heated at 60 °C for 30 min. Added the mix of click reagent gents (1 mM CuSO4, 100 µM BTTAA ligand and 100 µM biotin-PEG3 azide, 1 mM TCEP) and incubate for 60 min at RT. Protein samples were then precipitated with acetone and frozen at −20 °C overnight. Samples were then centrifuged 3,500 x g for 5 min at 4°C, aqueous/acetone solution was removed and protein precipitates and washed 3x with (cold acetone 1 mL, sonication and re-precipitated at −20 °C for 15 min, then spun at 3500 x g for 5 min at 4 °C). After the final wash, the pellets were air dried at RT for 10 min and re-solubilized in 100 µL 4% SDS PBS by sonication and gentle heat, then added PBS (2 mL) and diluted to 0.2% SDS. They were then enriched with streptavidin beads (100 µL per sample), incubated at room temperature for 1 h. Beads were then pelleted by centrifugation (1400 x g, 2 min), washed with 1% SDS PBS x 3, 2 M Urea PBS x 3, PBS X 3 each 2 mL). These samples were then split for immunoblot and proteomic analysis.

#### Tandem mass tag labeling for mass spectrometry

To the solution of the probe-bound streptavidin beads from above, ammonium bicarbonate (ABC) (25 mM) and 10 mM DTT 20 µL (100 mM stock) was added, then placed in 42 °C heat block for 30 min. 200 µL of 20 mM iodoacetamide was added and allowed to react at 37 °C for 30 min (protected from light). It was then spun down and washed with 10 mM DTT in 200 µL ABC, followed by washed 3 x with 25 mM ABC. The beads were pelleted by centrifugation (1,300 x g, 2 min) and resuspended in 200 µL of 25 mM ABC, 1mM CaCl2 2 µL, and trypsin 2 µg. The digestion was allowed to proceed overnight at 37 °C with shaking. Digested peptides were collected and dried using Genevac EZ- 2 evaporator. TMTsixplex^TM^ Isobaric Label Reagents (ThermoFisher Scientific), 0.8 mg per label, were equilibrated to room temperature, dissolved in 60 µL of dry acetonitrile and mixed by vortexing briefly before use. 7.5 µL of each TMT label reagent was carefully added to each sample (126 and 127 for the blank control, 128 and 129 for the probe, 130 and 131 for the competition) and incubated at room temperature for 1 h. 3 µL of 5% hydroxylamine was then added to each sample and incubated for 15 mins to quench the labeling reaction. Samples were then combined in equal quantities (about 100 µg), purified using the High pH Reversed-Phase Peptide Fractionation Kit (Pierce), and divided into two fractions (CQ83TMT1 and CQ83TMT2), and dried with a Genevac EZ-2 evaporator.

#### Tandem LC-MS/MS/MS

Mass spectrometry data was collected on an Orbitrap Fusion Lumos mass spectrometer coupled to an Easy-nLC 1200 (Thermo Fisher Scientific). Peptides were separated over a 180 min gradient of 0 to 50% acetonitrile in water with 0.1% formic acid at a flow rate of 300 nL/min on a 50 cm long PepMap RSLC C18 column (2 mm, 100 Å, 75 µm, x 50 cm). The full MS spectra were acquired in the Orbitrap at a resolution of 120,000. The 10 most intense MS1 ions were selected for MS2 analysis. The isolation width was set at 0.7 m/z and isolated precursors were fragmented by CID (35% CE). Following acquisition of each MS2 spectrum, a synchronous precursor selection (SPS) MS3 scan was collected on the top 10 most intense ions in the MS2 spectrum. The isolation width was set at 1.2 m/z and isolated precursors were fragmented using HCD. The mass spectrometry proteomics data will be made available at the ProteomeXchange Consortium (http://proteomecentral.proteomexchange.org) via the PRIDE partner repository (Perez-Riverol et al., 2019).

#### Proteomic analysis

MS raw files were analyzed using MaxQuant v1.6.17.0 by searching against the Uniprot human database supplemented with common contaminant protein sequences and quantifying according to SPS MS3 reporter ions. MaxQuant was run using the following parameters: reporter Ion MS3 – 6plex TMT; variable modifications - methionine oxidation (+15.995 Da), N-terminal protein acetylation (+42.011 Da), asparagine or glutamine deamidation (+0.984 Da); fixed modification - carbamido- methylation (+57.021 Da) of cysteine; digestion – trypsin/P.

#### IL-1β ELISA assays

2 × 10^5^ N/TERT-1 cells were plate on 6-well tissue culture plates and adhered overnight. Cells were then treated with compounds for the indicated time points and spent media samples were collected and clarified by centrifugation at 1000 x g for 1 min. Supernatants were harvested for quantitation using the R&D Human IL-1β quantikine ELISA kit according to the manufacturer’s instructions.

#### RNA-seq

2.5 × 10^6^ *CASP1*^−/−^ MV4;11 cells were plated in 6-well plates and treated with DMSO, MeBs (10 µM) or BFA (1.78 µM) in quadruplets for 6 h. Cellular RNA was extracted using the RNeasy Mini Kit (Qiagen) and residual genomic DNA was removed by on-membrane DNase digestion (Qiagen). The total RNA samples were sent to GENEWIZ for PolyA selection, library preparation, barcoding and sequencing using HiSeq configured for 2×150 bp paired-end reads (Illumina). An average of 28.9 × 10^6^ paired reads was generated per sample with a mean quality score of 35.81. The sequence reads were mapped to the *Homo sapiens* GRCh38 reference genome using STAR aligner v.2.5.2b, and unique gene hits were calculated using the Subread package v.1.5.2. The gene hits tables were then used for differential expression analysis by DESeq2. The Wald test was used to generate p-values and log2 fold changes.

#### Reverse Transcription – Quantitative Real-Time PCR (RT-qPCR)

Total RNA was isolated from HEK 293T or *CASP1*^−/−^ MV4;11 cells at the end of the experiment using the RNeasy Mini Kit (Qiagen) and reverse transcription-PCR was performed on 0.8 µg of mRNA using High Capacity cDNA Reverse Transcription Kit (Applied Biosystems). qPCR was performed on the cDNA for *CHAC1*, *DDIT3*, *DDIT4*, *SESN2*, *TRIB3*, and the housekeeping gene *GAPDH* using the indicated primer pairs and the PowerUp SYBR Green Master Mix dye (Applied Biosystems) on a QuantStudio 5 Real-Time PCR system (ThermoFisher). Data were analyzed using the ΔΔCt method in which ΔCt is the difference between the Ct value of the Gene of Interest and GAPDH, ΔΔCt is the difference between the treatment condition and DMSO, and the fold change is 2^−ΔΔCt^.

#### Puromycin cellular translation rate assay

HEK 293T cells were treated with the indicated compounds for 6 h prior to lysis. Lysates were treated with 10 µg/mL puromycin for 10 min, then immunoblot analysis was performed using an antibody raised against puromycin.

#### dTAG-CARD8 Assay

HEK293T cells stably expressing CASP1 and GSDMD were seeded at 1.25 × 10^5^ cells per well in 12-well tissue culture dishes. After 48 h, the cells were transfected with plasmids encoding dTAG-CARD8-ZUC (0.5 μg), CARD8 FIIND-S297A (0.3 μg) and RFP (0.2 μg) with FuGENE HD, according to the manufacturer’s instructions (Promega). At 24 h, cells were treated with DMSO, dTAG-13 (500 nM) and/or VbP (10 μM) for 3 h. Lysates were collected and protein content was evaluated by immunoblotting.

#### Statistical analysis

Two-sided Student’s t tests were used for significance testing unless stated otherwise. *P* values less than 0.05 were considered to be significant. Graphs and error bars represent means ± SEM of three independent experiments, unless stated otherwise. The investigators were not blinded in all experiments. All statistical analysis was performed using Microsoft Excel and GraphPad Prism 9.

### Experimental Procedures and Spectroscopic Data of Compounds

#### General Procedures

All reactions were carried out under an argon atmosphere with dry solvents under anhydrous conditions, unless otherwise noted. Reagents were purchased from Aldrich, Acros, or Fisher at the highest commercial quality and used without further purification, unless otherwise stated. Reactions were monitored by thin layer chromatography (TLC) carried out on MilliporeSigma glass TLC plates (silica gel 60 coated with F254, 250 μm) using UV light for visualization and aqueous ammonium cerium nitrate/ammonium molybdate or basic aqueous potassium permanganate as developing agent. NMR spectra were recorded on a Bruker Avance III 600 MHz. The spectra were calibrated by using residual undeuterated solvents (for ^1^H NMR) and deuterated solvents (for ^13^C NMR) as internal references: undeuterated chloroform (δH = 7.26 ppm) and CDCl3 (δC = 77.16 ppm); undeuterated methanol (δH = 3.31 ppm) and methanol-d4 (δC = 49.00 ppm). The following abbreviations are used to designate multiplicities: s = singlet, d = doublet, t = triplet, q = quartet, m = multiplet, br = broad. High-resolution mass spectra (HRMS) were recorded on a Waters Micromass LCT Premier XE TOF LC-MS. Scheme S1. Synthesis of CQ83 (**6**)

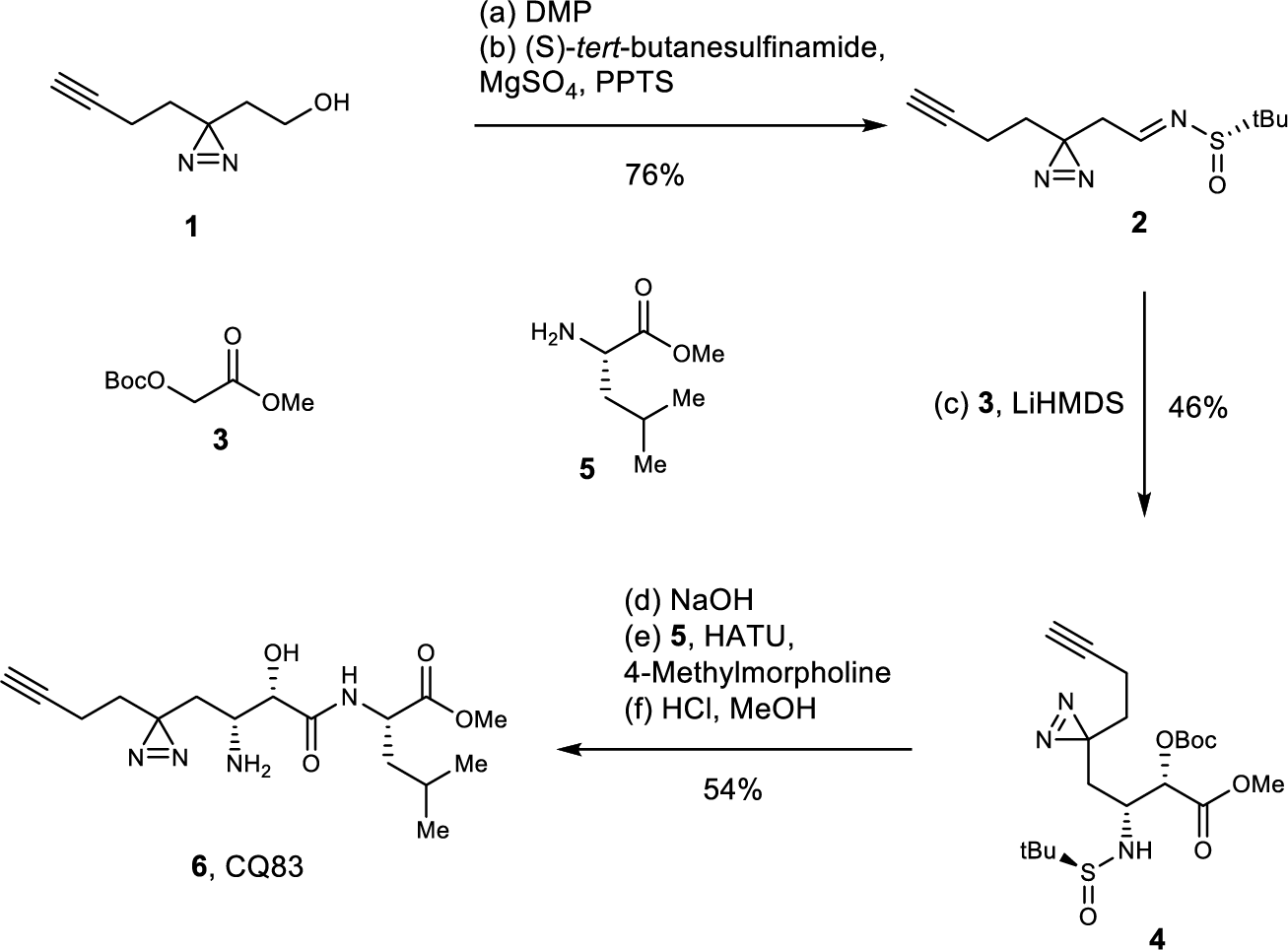

#### (S)-N-(2-(3-(but-3-yn-1-yl)-3*H*-diazirin-3-yl)ethylidene)-*tert*-butylsulfinamide (3)

To a solution of 2-(3-but-3-yn-1-yl-3*H*-diazirin-3-yl)-ethanol (50.0 mg, 0.362 mmol) in dry CH2Cl2 (1.2 mL) was added DMP (230 mg, 0.542 mmol) at 22 °C. The reaction mixture was stirred for 10 min at the same temperature before being passed through a short plug of silica gel with EtOAc/hexane (1:4) to give a colorless oil which was used for the next step without further purifications. To a solution of the crude aldehyde in dry CH2Cl2 (5 mL) were added pyridinium *p*-toluenesulfonate (PPTS, 4.8 mg, 0.0191 mmol), anhydrous MgSO4 (218 mg, 1.81 mmol) and (S)-*tert*- butanesulfinamide (66.0 mg, 0.545 mmol). The reaction mixture was stirred for 14 h at 22 °C. Then filtered through a pad of celite, washed with CH2Cl2 and concentrated under vacuum. The residue was passed through a short plug of silica gel with EtOAc/hexane (1:4) to give the desired imine **2** (66.1 mg, 76%) as a colorless oil. **2**: ^1^H NMR (600 MHz, CDCl3): δ = δ 8.01 (t, *J* = 4.4 Hz, 1 H), 2.58 (dd, *J* = 17.0, 4.4 Hz, 1 H), 2.52 (dd, *J* = 17.1, 4.3 Hz, 1 H), 2.06 (ddd, *J* = 7.2, 2.8, 0.8 Hz, 1 H), 2.04 (dd, *J* = 7.2, 2.7 Hz, 1 H), 2.01 (t, *J* = 2.7 Hz, 1 H), 1.82 – 1.71 (m, 2 H), 1.24 (s, 9 H) ppm; ^13^C NMR (151 MHz, CDCl3): δ = 163.5, 82.5, 69.8, 57.2, 40.8, 32.3, 26.1, 22.6, 13.4 ppm; HRMS (*m/z*): [M+H]^+^ calcd for C11H18N3OS^+^ 240.1171, found 240.1171.

#### Methyl (2S,3R)-4-(3-(but-3-yn-1-yl)-3*H*-diazirin-3-yl)-2-((*tert*-butoxycarbonyl)oxy)-3-(((S)- *tert*-butylsulfinyl)amino)butanoate (4)

A solution of methyl 2-((*tert*-butoxycarbonyl)oxy)acetate **3** (213 mg, 1.12 mmol) in dry THF (3 mL) maintained under an atmosphere of argon was cooled to –78 °C and then treated with LiHMDS (1.12 mL, 1.0 M solution in THF, 1.12 mmol). The reaction mixture was stirred for 1 h at the same temperature before imine **2** (53 mg, 0.221 mmol) in THF (200 uL) was added slowly. The mixture was allowed to stir for 5 h before it was quenched with saturated aq. NH4Cl (5 mL). The aqueous phase was extracted with EtOAc (3 × 10 mL). The combined organic phases were washed with brine (10 mL), dried over anhydrous MgSO4, filtered and concentrated under vacuum. The residue was passed through a short plug of silica gel with EtOAc/hexane (1:3) to give the desired methyl ester **4** (43.7 mg, 46%) as a colorless oil. **4**: ^1^H NMR (600 MHz, CDCl3): δ = 4.91 (br s, 1 H), 4.45 (dd, *J* = 11.8, 5.8 Hz, 1 H), 3.79 (s, 3 H), 3.55 (ddd, *J* = 10.8, 5.9, 3.6 Hz, 1 H), 2.85 (br s, 1 H), 2.07 – 1.88 (m, 4 H), 1.67 – 1.52 (m, 2 H), 1.50 (s, 9 H), 1.14 (s, 9 H) ppm; ^13^C NMR (151 MHz, CDCl3): δ = δ 172.9, 155.6, 84.6, 82.6, 77.4, 72.4, 69.5, 60.4, 52.4, 46.2, 32.4, 28.3, 26.1, 22.9, 13.4 ppm; HRMS (*m/z*): [M+H]^+^ calcd for C19H32N3O6S^+^ 430.2012, found 430.2002.

#### Methyl((2S,3R)-3-amino-4-(3-(but-3-yn-1-yl)-3*H*-diazirin-3-yl)-2-hydroxybutanoyl)-L- leucinate (6)

To the solution of methyl ester **3** (36 mg, 0.0838 mmol) in 1,4-dioxane/H2O (1:1, 1.0 mL was added NaOH (3.0 mg, 0.125 mmol) and the reaction mixture was stirred at 22 °C for 1 h. The mixture was acidified to pH 3-4 with Dowex® 50W X8 resin. The resin was filtered and washed with CH2Cl2. The aqueous phase was extracted with CH2Cl2 (3 × 10 mL). The combined organic phases were washed with brine (10 mL), dried over anhydrous MgSO4, filtered and concentrated under vacuum to give a colorless oil which was used for the next step without further purifications. To a solution of crude oil from the last step in CH2Cl2 (2.0 mL) were sequentially added L-Leucine methyl ester hydrochloride **5** (18.2 mg, 0.100 mmol), HATU (38.1 mg, 0.100 mmol), 4-Methylmorpholine (21.2 mg, 23 μL, 0.209 mmol, at 0 °C. The reaction mixture was allowed to stir for another 3 h before it was quenched by addition of saturated aq. NaHCO3 solution (2 mL). The organic layer was separated, and the aqueous layer was extracted with CH2Cl2 (3 × 5 mL). The organic layers were combined, washed with brine (10 mL), dried over Na2SO4, and concentrated under vacuum. The resulting residue was purified by flash column chromatography (silica gel, EtOAc:hexane = 1:4, *v*/*v* → 1:1, *v*/*v*) to give the desired amide as a colorless oil. To a stirred solution of the obtained oil in MeOH (0.3 mL) was added HCl (0.3 mL, 3.0 M solution in MeOH, 0.9 mmol) at 0 °C. The reaction mixture was warmed 22 °C and stirred for 24 h at the same temperature. The mixture was concentrated under vacuum, and the residue was purified by recrystallization from MeOH/diethyl ether to give **6** (17.0 mg, 54% for 3 steps) as a white solid. **6**: ^1^H NMR (600 MHz, methanol-d4): δ = 4.47 (dd, *J* = 9.6, 4.5 Hz, 1 H), 4.33 (d, *J* = 3.5 Hz, 1 H), 3.74 (s, 3 H), 3.51 (dd, *J* = 6.9, 3.4 Hz, 1 H), 2.34 (d, *J* = 2.7 Hz, 1 H), 2.08 – 2.03 (m, 2 H), 1.96 (dd, *J* = 15.3, 7.1 Hz, 1 H), 1.77 (dq, *J* = 15.6, 8.3, 7.8 Hz, 1 H), 1.73 – 1.65 (m, 5 H), 0.98 (d, *J* = 5.6 Hz, 3 H), 0.95 (d, *J* = 5.6 Hz, 3 H) ppm; ^13^C NMR (151 MHz, methanol-d4): δ = 174.4, 173.1, 83.3, 70.8, 70.5, 52.9, 52.4, 50.9, 41.1, 34.3, 32.7, 26.5, 26.0, 23.2, 22.0, 13.7. ppm; HRMS (*m/z*): [M+H]^+^ calcd for C16H27N4O4^+^ 339.2032, found 339.2022.

