## Supplemental Figures 1-7, Tables 1-2 for "Cytosolic peptide accumulation activates the NLRP1 and CARD8 inflammasomes"

#### SUPPLEMENTARY INFORMATION

### Figure S1. AP inhibition enhances DPP8/9 inhibitor-induced cell death, related to Figure 2.

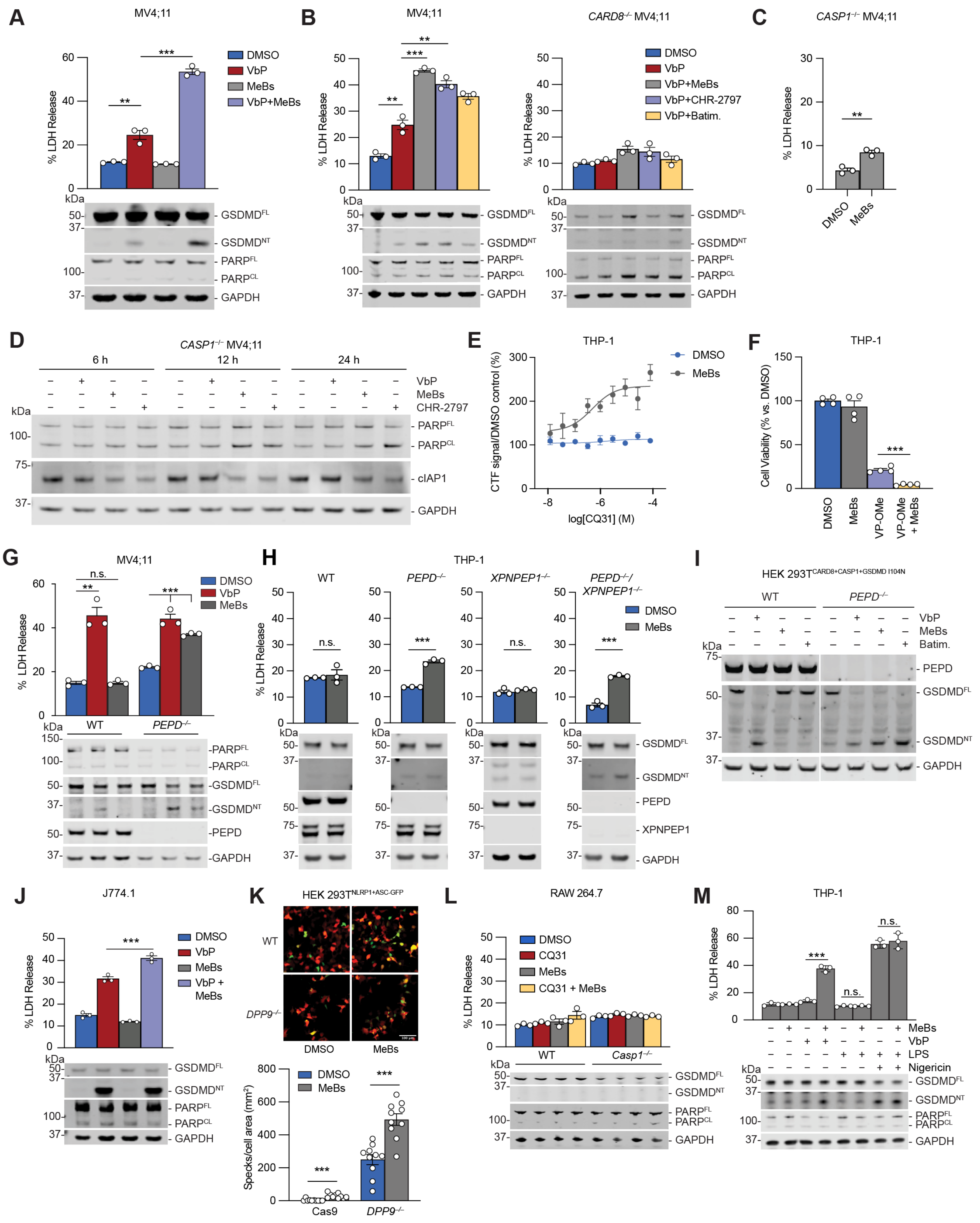

**Figure S1. AP inhibition enhances DPP8/9 inhibitor-induced cell death, related to Figure 2.** (**A,B**) WT or *CARD8*$^{-/-}$ MV4;11 cells were treated with aminopeptidase inhibitors VbP (10 μM), MeBs (10 μM), batimastat (Batim.; 10 μM), and/or CHR-2797 (10 μM) for 6 h before LDH release and immunoblot analyses. (**C**) *CASP1*$^{-/-}$ MV4;11 cells were treated with MeBs (10 μM) for 24 h prior to cell viability assessment by LDH release. (**D**) *CASP1*$^{-/-}$ MV4;11 cells were treated with VbP (10 μM), MeBs (10 μM) or CHR-2797 (10 μM) for the indicated amount of time before immunoblot analysis. (**E,F**) THP-1 cells were treated with MeBs (10 μM), VP-OMe (10 μM), and/or the indicated concentration of CQ31 for 6 h. Cytotoxicity was then assessed by CTF (**E**) or CTG (**F**). Data are means ± SEM of 4 replicates (**E,F**). (**G**) WT or *PEPD*$^{-/-}$ MV4;11 cells were treated with VbP (2 μM) or MeBs (1 μM) for 24 h before LDH release and immunoblot analyses. (**H**) WT, *PEPD*$^{-/-}$, *XPNPEP1*$^{-/-}$, or *PEPD*$^{-/-}$/*XPNPEP1*$^{-/-}$ THP-1 cells were treated with MeBs (10 μM) for 6 h before LDH release and immunoblot analyses. (**I**) WT or *PEPD*$^{-/-}$ HEK 293T cells ectopically expressing CARD8, CASP1 and GSDMD-I104N (a defective pore-forming mutant) were treated with VbP (10 μM), MeBs (10 μM) or Batim. (10 μM) for 6 h before immunoblot analysis. (**J**) J774.1 cells were treated with VbP (2 μM) and/or MeBs (2 μM) for 6 h followed by LDH release and immunoblot analyses. (**K**) WT or *DPP9*$^{-/-}$ HEK 293T cells ectopically expressing NLRP1 and ASC-GFP were treated with DMSO or MeBs (20 μM) for 6 h. ASC speck formation was assessed by fluorescence microscopy. Representative images and average cells with specks (%) ± SEM of 10 replicates are shown. (**L**) RAW 264.7 WT or *Casp1*$^{-/-}$ cells were treated with CQ31 (20 μM) and/or MeBs (10 μM) for 24 h before LDH and immunoblot analyses. (**M**) THP-1 cells were primed with LPS (5 μg/mL) for 18 h followed by treatment with MeBs (10 μM, 6 h), VbP (10 μM, 6 h) and/or nigericin (10 μM, 30 min) before LDH release and immunoblot analyses. Data are means ± SEM of 3 replicates unless otherwise stated. All data, including immunoblots, are representative of three or more independent experiments. *** $p < 0.001$, ** $p < 0.01$ by two-sided Students *t*-test. n.s., not significant.

### Figure S2. MeBs accelerates NT degradation, related to Figure 3.

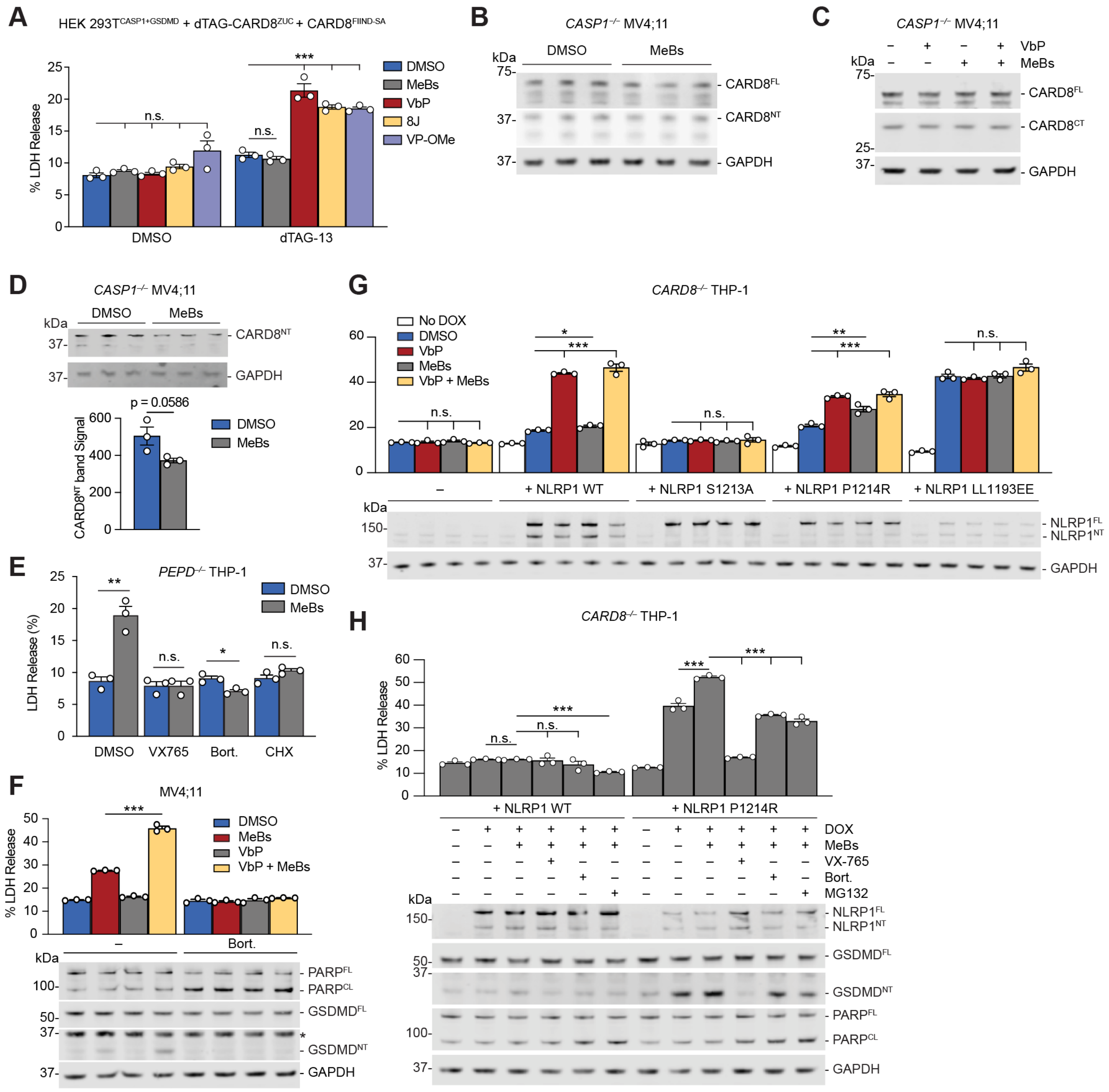

**Figure S2. MeBs accelerates NT degradation, related to Figure 3.** (**A**) HEK 293T$^{CASP1+GSDMD}$ cells were transiently transfected plasmids encoding dTAG-CARD8$^{ZUC}$ and the isolated FIIND$^{SA}$ 48 h prior to treatment with dTAG-13 (500 nM), MeBs (10 μM), VbP (10 μM), compound 8j (50 μM) or valine-proline methyl ester (VP-OMe, 0.5 mM) for 6 h. Supernatants were then assessed for LDH release. (**B-D**) *CASP1*$^{-/-}$ MV4;11 cells were treated with VbP (10 μM) and/or MeBs (10 μM), or the combination for 6 h (**B,C**) or 48 h (**D**) followed by immunoblot and/or densitometry analysis of images. (**E**) *PEPD*$^{-/-}$ THP-1 cells were treated with MeBs (10 μM), VX765 (50 μM) and/or Bort. (1 μM) for 8 h followed by LDH release analysis. (**F**) MV4;11 cells were treated with MeBs (10 mM), VbP (10 μM) and/or Bort. (10 μM) for 6 h followed by LDH release and immunoblot analyses. (**G**,**H**) *CARD8*$^{-/-}$ THP-1 cells stably expressing doxycycline (DOX) inducible NLRP1 WT, S1213A, P1214R, or LL1193EE were induced with 1 μg/mL DOX for 17 h (**G**) or 3 h (**H**) prior to treatment with VbP (10 μM), MeBs (10 μM), VX765 (50 μM), bort. (10 μM) and/or MG132 (10 μM) for 6 h. Cells death was then assessed using LDH release and immunoblot analyses. Data are means ± SEM of 3 replicates. All data, including immunoblots, are representative of three or more independent experiments. *** $p < 0.001$, ** $p < 0.01$, * $p < 0.05$ by two-sided Students *t*-test. n.s., not significant.

**Figure S3. Investigating the impact of AP inhibition on amino acid recycling, related to Figure 4.**

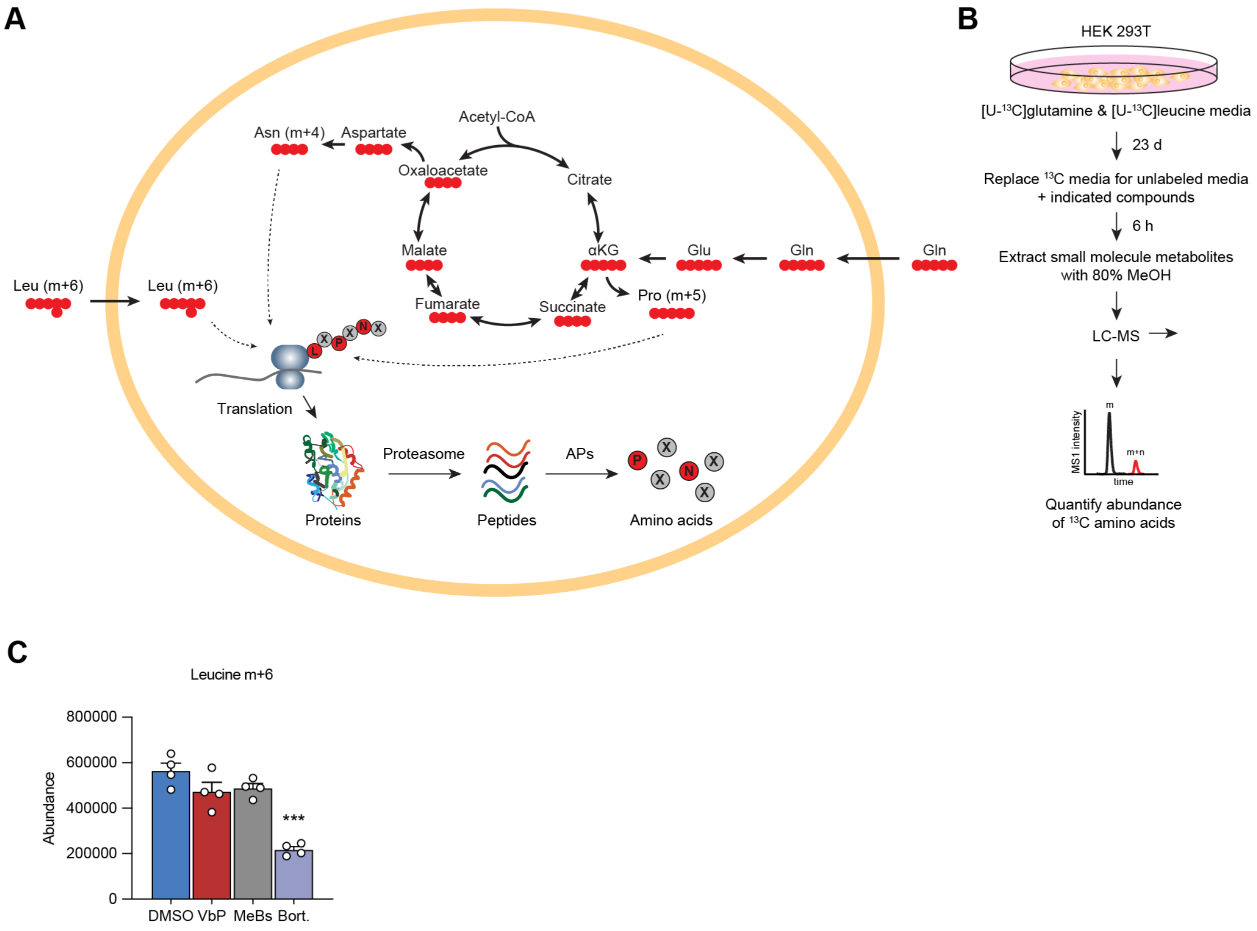


**Figure S3. Investigating the impact of AP inhibition on amino acid recycling, related to Figure 4.** (**A**) Schematic of amino acid stable isotope incorporation into proteins. (**B**) Overview of stable isotope tracing experiment. HEK 293T cells were grown for 3 weeks in [U-$^{13}$C] L-glutamine and [U-$^{13}$C] L-leucine labeled DMEM. Cells were plated in labeled media, and then next day media was simultaneously replaced with media containing unlabeled glutamine and leucine and treated with the indicated compounds for 6 h before metabolite extraction and isotope abundance measurement. (**C**) Abundance of m+6 labeled leucine in the isotope tracing experiment. This experiment was performed once. *** $p < 0.001$ by two-sided Students *t*-test.

**Figure S4. Differential breakdown of proline-containing peptides, related to Figure 5.**

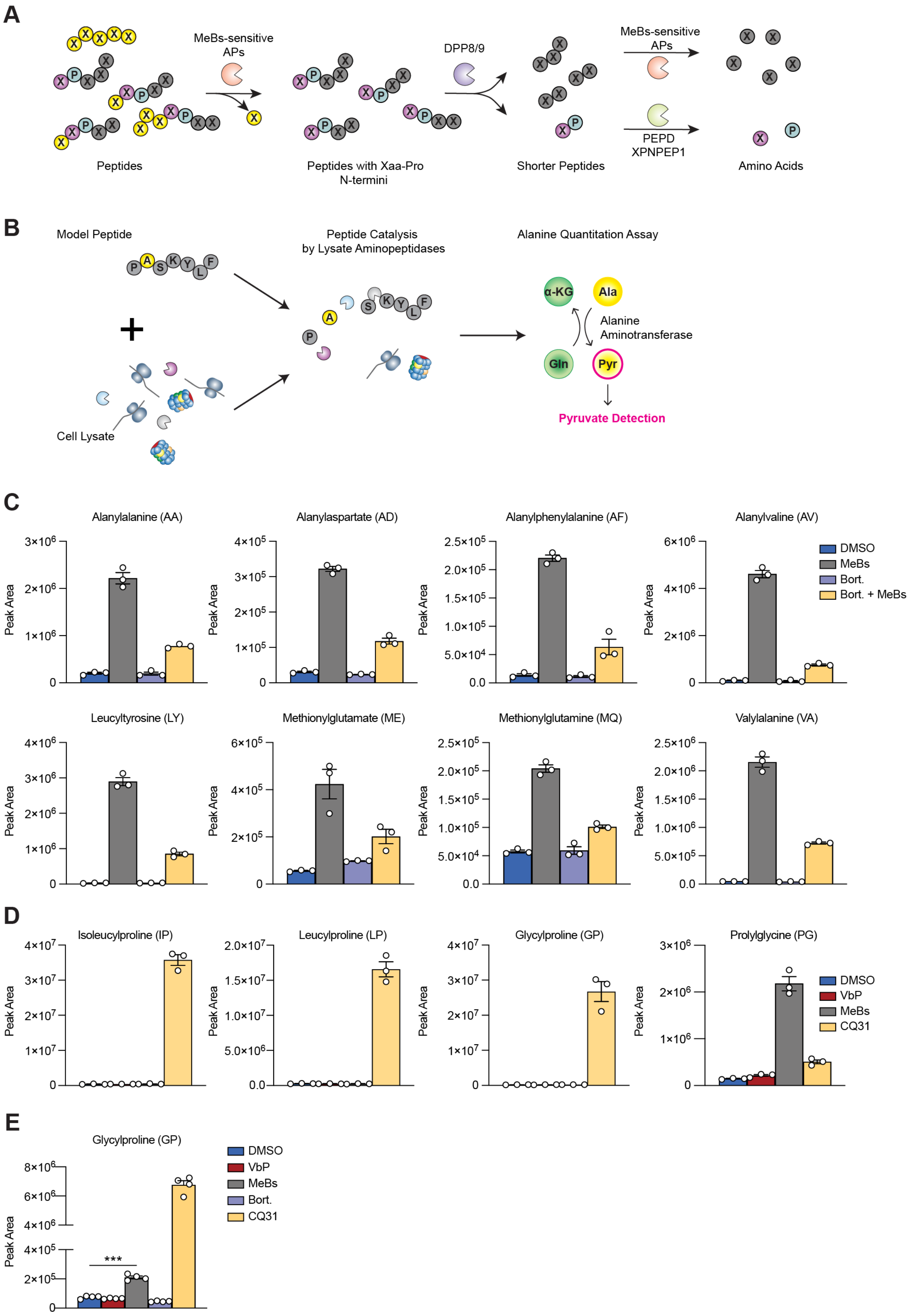

**Figure S4. Differential breakdown of proline-containing peptides, related to Figure 5**. (**A**) Overview of peptide breakdown by APs. APs typically remove amino acids sequentially one at a time from the N-termini of peptides (yellow amino acids). However, most APs (except XPNPEP1) cannot cleave before proline due proline's conformationally restricted bonds (purple amino acids). Thus, when proline is the second position from a peptide's N-terminus, DPP8/9 cleave the Xaa-Pro dipeptide, which is subsequently processed by M24B family aminopeptidases PEPD (and potentially XPNPEP1). The APs can then continue processing the peptide (gray amino acids). (**B**) Overview of model peptide cleavage assay. The peptide PASKYLF is mixed with HEK 293T cell lysates for 6 h, followed by measurement of peptide cleavage by an alanine detection assay. (**C**) THP-1 cells were pre-treated with DMSO or Bort. (10 μM) for 30 min, and then treated with DMSO or MeBs (10 μM) for 5.5 h. Intracellular metabolites were then extracted and dipeptide concentrations were measured by LC-MS. (**D**) HEK 293T cells were treated with MeBs (10 μM), VbP (10 μM), or CQ31 (20 μM) for 24 h. Intracellular metabolites were extracted, and the indicated dipeptide concentrations were measured by LC-MS. This experiment was previous incorporated in Rao et al., 2022, but the data with MeBs is original to this manuscript. (**E**) HEK 293T cells were treated with VbP (10 μM), MeBs (10 μM), or Bortezomib (10 μM) for 6 h. Intracellular metabolites were extracted and the relative abundance of Gly-Pro was measured by LC-MS. Data are means ± SEM of 3 or more replicates. All metabolomics data were performed in a single independent experiment. *** $p < 0.001$, ** $p < 0.01$, * $p < 0.05$ by two-sided Students *t*-test. n.s., not significant.

**Figure S5. AP inhibitors cause both cIAP1 depletion and pyroptosis, related to Figure 6.**

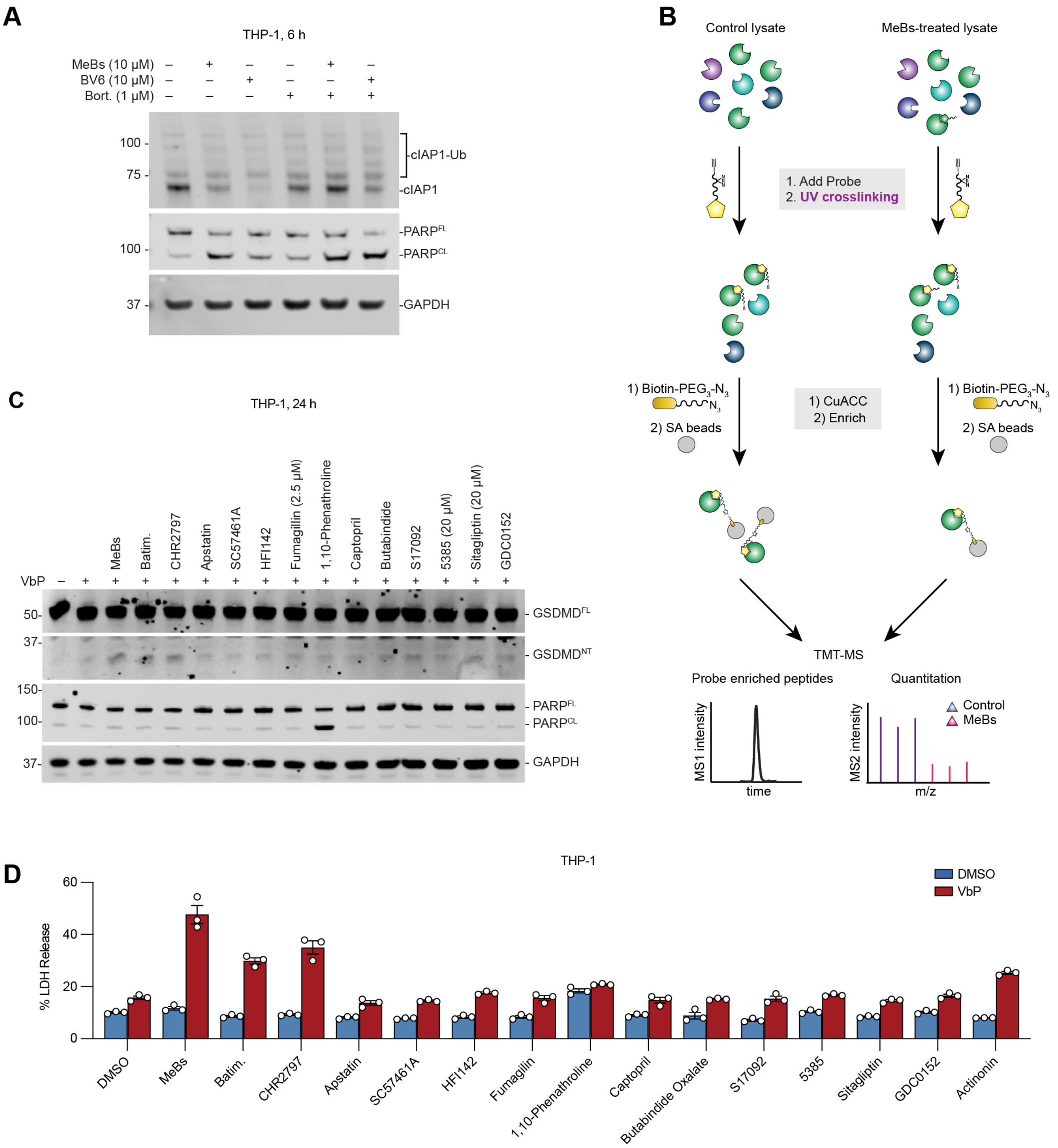


**Figure S5. AP inhibitors cause both cIAP1 depletion and pyroptosis, related to Figure 6.** (**A**) THP-1 cells treated with MeBs (10 µM), BV6 (10 µM) and/or Bort. (1 µM) for 6 h before immunoblotting analysis. (**B**) Schematic of target identification protocol using the CQ83 probe. (**C**,**D**) THP-1 cells treated with VbP (2 µM) and the indicated AP inhibitors (all 10 µM except for fumagillin, 2.5 µM; 5385, 20 µM; sitagliptin, 20 µM) or GDC-0152 (5 µM) for 18 h prior to immunoblotting (**C**) or LDH (**D**) analyses. Data are means ± SEM of 3 replicates. All data, including immunoblots, are representative of three or more independent experiments.

**Figure S6. Unfolded proteins cause degradation of NT fragments, related to Figure 7.**

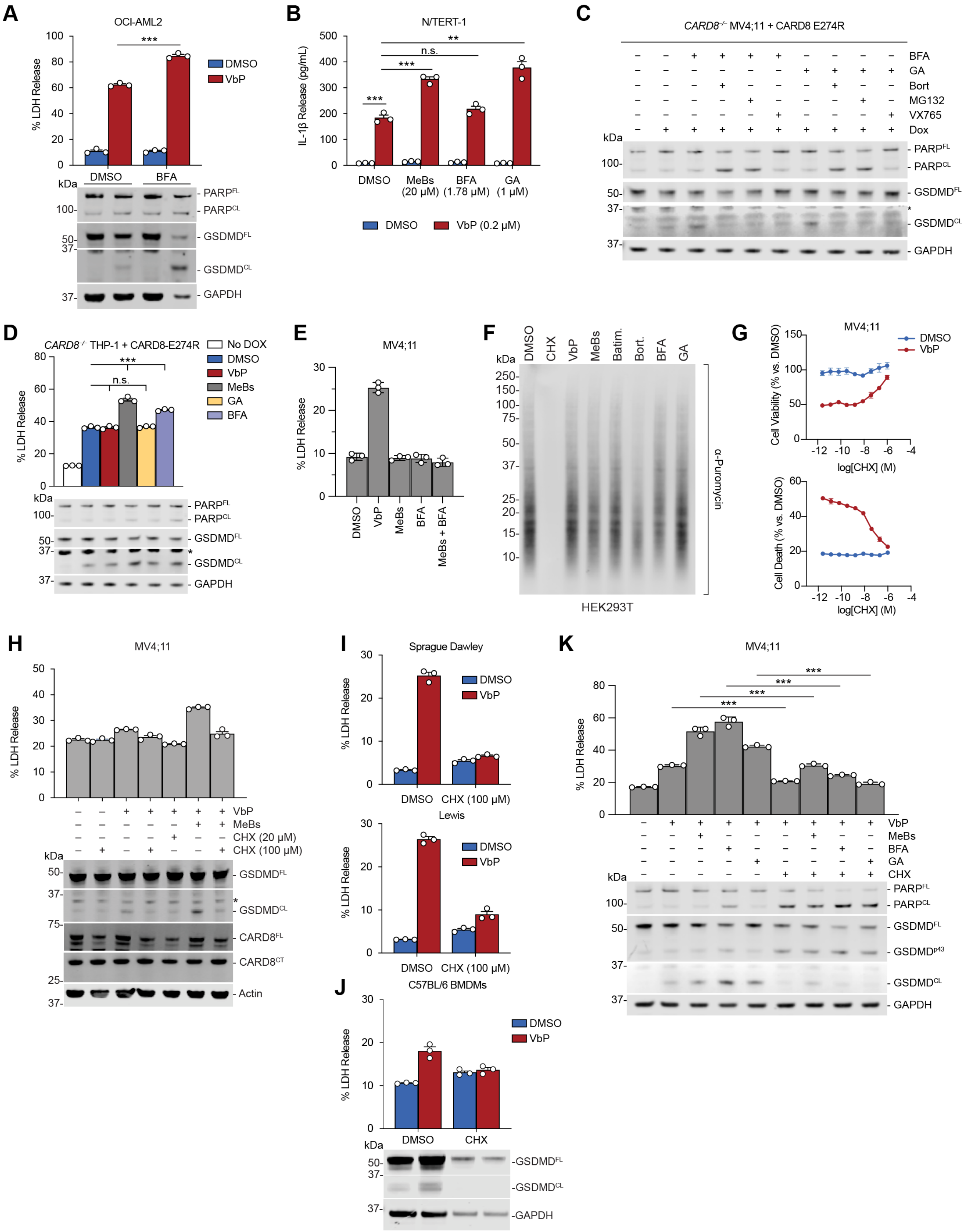

**Figure S6. Unfolded proteins cause degradation of NT fragments, related to Figure 7.** (**A**) OCI-AML2 cells were treated with VbP (10 μM) and/or BFA (1.78 μM) for 6 h prior to LDH release and immunoblot analyses. (**B**) N/TERT-1 human keratinocytes were treated with MeBs (20 μM), GA (1 μM) and/or VbP (0.2 μM) for 8 h prior to IL-1β release and immunoblot analyses. (**C**) *CARD8*$^{-/-}$ MV4;11 cells stably expressing doxycycline (DOX)-inducible CARD8 E274R were induced with 100 ng/mL DOX for 16 h prior to treatment with BFA (1.78 μM), GA (1 μM), Bortezomib (Bort., 10 μM), MG132 (10 μM) and/or VX765 (50 μM) for 6 h. Cell death was then evaluated by immunoblot analyses. (**D**) *CARD8*$^{-/-}$ THP-1 cells stably expressing doxycycline (DOX)-inducible CARD8 E274R were induced with 100 ng/mL DOX for 16 h prior to treatment with VbP (10 μM), MeBs (10 μM), GA (1 μM), and/or BFA (1.78 μM) for 6 h. Cell death was then evaluated by LDH release and immunoblot analyses. (**E**) MV4;11 cells were treated with VbP (10 μM), MeBs (10 μM) and/or BFA (1.78 μM) for 6 h. Cell death was then evaluated by LDH release. (**F**) HEK 293T cells were treated with CHX (20 μM), VbP (10 μM), MeBs (10 μM), Batim. (10 μM), Bort. (10 μM), BFA (1.78 μM) or GA (1 μM) for 6 h. Puromycin was then added to cells for 10 min to modify nascent proteins, which were detected by immunoblotting against puromycin. (**G,H**) MV4;11 cells were treated with VbP (10 μM), MeBs (10 μM) and/or the indicated concentration of CHX for 6 h before cell viability was assessed by CTG (**G,** *top*) or CTF (**G,** *bottom*) or LDH release and immunoblotting analyses (**H**). (**I**) Sprague Dawley (*top*) or Lewis (*bottom*) rat bone marrow derived macrophages were treated with VbP (10 μM) and/or CHX (50 μM) for 6 h. Cell death was then evaluated by LDH release analysis. (**J**) C57BL/6 mouse bone marrow derived macrophages (BMDMs) were treated with VbP (2 μM) and/or CHX (100 μM) for 6 h. Cell death was then evaluated by LDH release and immunoblot analyses. (**K**) MV4;11 cells were treated with VbP (10 μM), MeBs (10 μM), GA (1 μM), BFA (1.78 μM), and/or CHX (100 μM) for 6 h. Cell death was then evaluated by LDH release and immunoblot analyses. Data are means ± SEM of 3 or more replicates. All data, including immunoblots, are representative of three or more independent experiments. *** $p < 0.001$, ** $p < 0.01$ by two-sided Students *t*-test. n.s., not significant.

**Figure S7. Peptides activate the CARD8 and NLRP1 inflammasomes.**

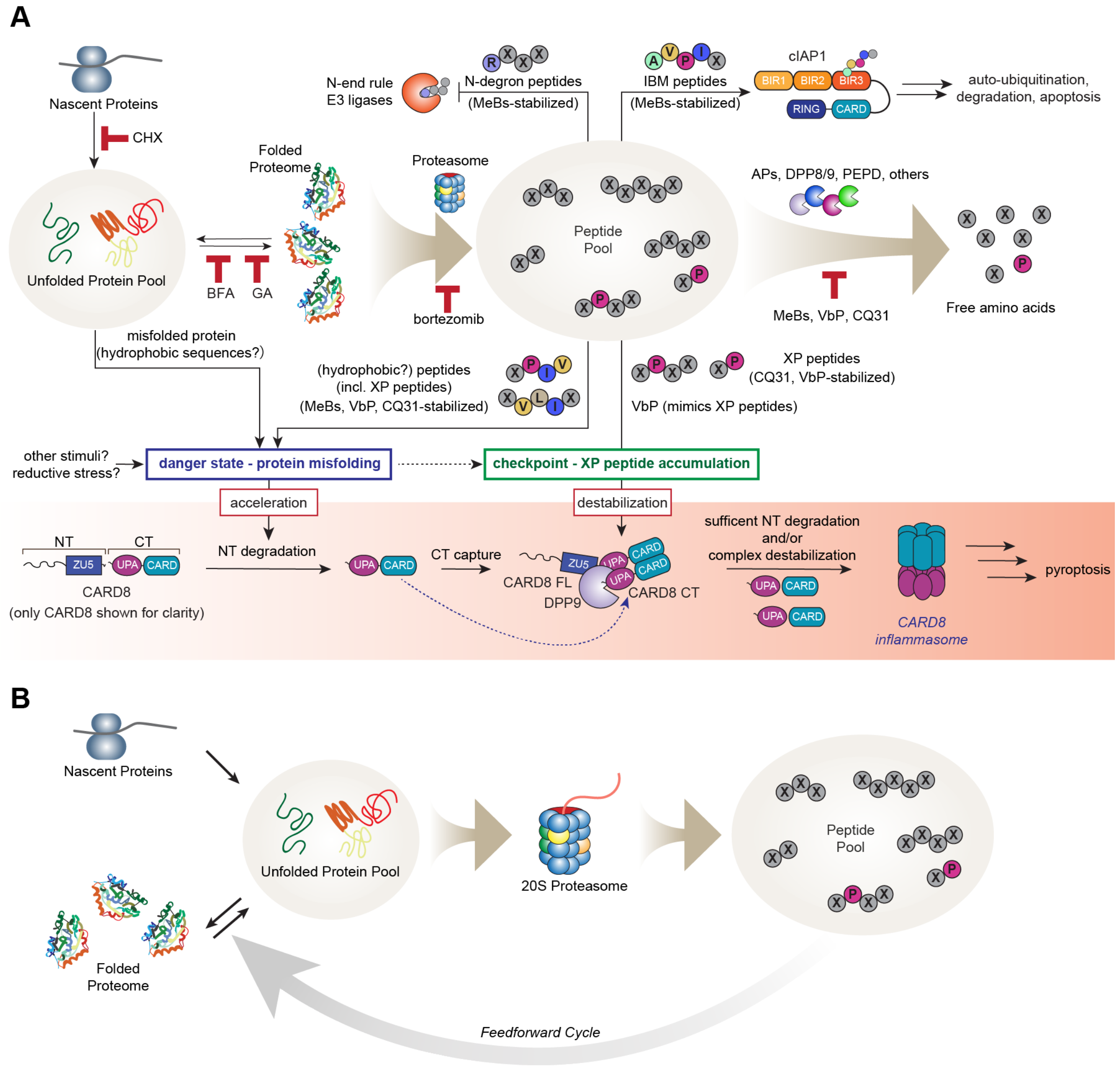


**Figure S7. Peptides activate the CARD8 and NLRP1 inflammasomes.** (**A**) Agents that interfere with protein folding, including peptides, accelerate NT degradation. Peptides with XP N-termini also destabilize the repressive ternary complexes. Peptides with distinct N-terminal sequences interfere with other cellular processes, including the N-end rule and IAP pathways. (**B**) Peptide accumulation can interfere with protein folding. Unfolded proteins are rapidly degraded, thereby releasing even more peptides. These peptides, if not quickly destroyed, could in turn similarly interfere with protein folding in a feedforward cycle of proteotoxic stress.

**Table S1. Sensitivities of CARD8 and NLRP1 alleles to various stimuli, related to Figure 1.** mNLRP1B alleles 3 and 4 are not shown because they are non-functional inflammasome sensors. IpaH7.8 is a *Shigella flexneri* E3 ligase. NT, not tested.

| | | Sensitivity | | | | |
|---|---|---|---|---|---|---|
| **Species** | **Inflammasome sensor** | **VbP** | **CQ31** | **dsRNA** | **IpaH7.8** | **Protease** |
| human | CARD8 | Yes | Yes | No | NT | HIV-1 protease |
| human | NLRP1 | Yes | No | Yes | No | Enteroviral 3C proteases |
| mouse | NLRP1B - allele 1 | Yes | No | No | Yes | Lethal factor |
| mouse | NLRP1B - allele 2 | Yes | No | NT | No | Enteroviral 3C proteases |
| mouse | NLRP1B - allele 5 | Yes | NT | NT | NT | Lethal factor |
| mouse | NLRP1A | Yes | No | NT | NT | none known |
| rat | NLRP1 - allele 1 | Yes | NT | NT | NT | Lethal factor |
| rat | NLRP1 - allele 2 | Yes | NT | NT | NT | Lethal factor |
| rat | NLRP1 - allele 3 | Yes | NT | NT | NT | none known |
| rat | NLRP1 - allele 4 | Yes | NT | NT | NT | none known |
| rat | NLRP1 - allele 5 | Yes | NT | NT | NT | none known |

**Table S2. Summary of CARD8 and NLRP1 alleles in various biological systems, related to Figure 1.** The sensitivity of each experimental system to the combination of VbP and MeBs is shown. NT, not tested.

| | | | | VbP + MeBs Synergy | |
|---|---|---|---|---|---|
| **Identifier** | **Type** | **Species** | **Sensor** | **Prior work** | **This study** |
| THP-1 | cell line | human | CARD8 | Yes | Yes |
| MV4;11 | cell line | human | CARD8 | NT | Yes |
| OCI-AML2 | cell line | human | CARD8 | NT | Yes |
| CD3 T cells | primary cells | human | CARD8 | NT | Yes |
| RAW264.7 | cell line | mouse | NLRP1B-allele 1 | Yes | Yes |
| J774.1 | cell line | mouse | NLRP1B-allele 1 | NT | Yes |
| N/TERT-1 | cell line | human | NLRP1 | NT | Yes |
| C57BL/6J mouse | animal | mouse | NLRP1B-allele 2; NLRP1A | Yes | NT |
| Sprague Dawley BMDMs | primary cells | rat | NLRP1-allele 1 | Yes | NT |

**Table S3. Table of amino acids concentrations, related to Figure 4.**

**Table S4. List of proteins enriched by CQ83 analyzed by TMT-mass spectrometry, related to Figure 6.**

**Table S5. Table of transcript counts measured by RNA-Seq, related to Figure 7.**
